# Representation of multiple objects in macaque category-selective areas

**DOI:** 10.1101/264465

**Authors:** Pinglei Bao, Doris Y. Tsao

## Abstract

Object recognition in the natural world usually occurs in the presence of multiple surrounding objects, but responses of neurons in inferotemporal (IT) cortex, the large brain area responsible for object recognition, have mostly been studied only to isolated objects. We study rules governing responses to multiple objects by cells in two category-selective regions of macaque IT cortex, the middle lateral face patch (ML) and the middle body patch (MB). We find that responses of single ML and MB cells to pairs of objects could be explained by the widely-accepted framework of normalization, with one added ingredient: homogeneous category selectivity of neighboring neurons forming the normalization pool. This rule leads to winner-take-all, contralateral-take-all, or weighted averaging behavior in single cells, depending on the category, spatial configuration, and relative contrast of the two objects. The winner-take-all behavior suggests a potential mechanism for clutter-invariant representation of face and bodies under certain conditions.

## Introduction

In the real world, primates need to recognize objects in the presence of other objects, since objects seldom appear in isolation. Behavioral evidence suggests that both humans and macaque monkeys are able to do this readily^1, 2^. What is the neural mechanism underlying representation of multiple objects?

One simple notion is that the representation of objects in inferotemporal cortex (IT), the end stage of the ventral visual partway, should be invariant to the presence of other objects, i.e., a neuron’s response to its preferred object should not be different when the object is presented alone compared to when it is presented with other objects. In other words, cells should implement a “winner-take-all” rule, responding to a collection of objects as if only the most preferred object were present. This would be a highly non-trivial computation: IT cells have large receptive fields encompassing multiple objects, and a winner-take-all rule would require some way to shut off inputs representing the non-preferred object. However, most previous electrophysiological studies of multiple object representation in IT find that the responses of cells are not invariant to the presence of clutter, i.e., cells are not implementing a winner-take-all rule. Sheinberg and Logothetis trained monkeys to look for a target object in a cluttered background, and found that IT neurons showed bursts shortly before effective targets were fixated, but the magnitude of these bursts was often smaller than that to isolated targets^3^. Many other studies have reported weaker responses in IT to object pairs compared to isolated, preferred objects^4, 5, 6, 7^, consistent with findings in early visual cortex ^8^.

In particular, it has been claimed that an extremely simple rule can describe the response of most IT cells to multiple objects: averaging of the responses to the individual objects, regardless whether the objects are preferred or non-preferred^4^. Computational simulations show that an averaging rule can permit limited clutter-invariant recognition through population coding^9^.

However, recognition performance is significantly worse than with a winner-take-all rule^10^. Thus many researchers assume that top-down attention provides the brain’s primary solution to visual recognition in clutter^11, 12^.

However, previous electrophysiological studies exploring the rules governing responses to multiple objects (a process we refer to as “multiple object integration” below) in IT during passive fixation suffered one important limitation: they all recorded from randomly selected IT neurons whose role in coding the object set tested is unknown^4, 6, 7, 13^. Up to now, this limitation has been difficult to overcome: for most cells in IT, the only clue we have to whether the cell is involved in encoding a particular object is whether the cell under study responds to the object. But category-selective regions of IT cortex provide an exception to this rule. For example, multiple lines of evidence suggest that the macaque face patch sysyem is specialized for coding faces^14, 15, 16^ and the macaque body patch system is specialized for coding bodies^17, 18, 19^. Thus the rules used by cells in face/body patches for multiple object integration involving faces/bodies have higher likelihood to be behaviorally relevant than those used by randomly sampled IT cells for multiple object integration involving random objects.

Indeed, in contrast to macaque electrophysiology studies, several human fMRI studies have explored the question of how the brain processes multiple objects specifically within category-selective regions and found evidence for clutter-tolerant representation of the preferred category in these regions. Decoding of object category from multivoxel fMRI response patterns in face and place-selective areas is more tolerant to clutter than decoding in non-category selective IT regions^20, 21^. Furthermore, face and body detection is highly efficient even in cluttered displays^22,23^, and behavioral performance on a change detection task in a multiple object display is superior when objects are drawn from categories represented by distinct category-selective regions^24^, further suggesting that regions selective for a particular category can filter out representations of objects from other categories. Thus there is a discrepancy between human fMRI studies and macaque single-unit studies, with respect to the mechanism for multiple object representation in IT cortex. To resolve this discrepancy, it is essential to obtain single-cell data from fMRI-identified category-selective areas.

In the present study, we re-investigated the question of how cells in IT cortex respond to multiple objects through targeted recordings in face and body patches. We targeted neurons in the middle lateral face patch (ML) of three monkeys and the middle body patch (MB) of two monkeys and studied responses to multiple object stimuli in a passive fixation paradigm. The rules for integrating preferred and non-preferred stimuli in ML and MB turned out to be very different from a simple averaging rule proposed previously based on recordings in randomly selected IT neurons^4, 7^. We found that single ML and MB cells could switch between one of three different behaviors, winner-take-all, contralateral-take-all, or weighted averaging, depending on the category, spatial configuration, and relative contrast of the two objects. The finding of winner-take-all and contralateral-take-all behavior in face and body patches suggests a new mechanism by which clutter invariance can be solved. Furthermore, the category-dependent integration behavior observed in the face and body patches underscores the importance of studying integration mechanisms in a manner that respects the functional architecture of IT. We show how our results arise naturally from the widely-accepted framework of normalization^25^, with one added ingredient: homogenous category selectivity of neighboring neurons forming the normalization pool.

## Results

### Localization of face patches with fMRI

We localized face patches in three monkeys with fMRI by presenting a face localizer stimulus set containing images of faces and non-face objects^14, 26^, and targeted middle face patch ML for electrophysiological recording. Face-selective units were first identified by presenting 16 faces and 80 non-face objects in the fovea. Consistent with previous studies, we found 90% of cells had a face selectivity index greater than 0.33 (Supplementary Figure 1, see Methods for details); these units were selected for further study.

### Response to a preferred and a non-preferred stimulus in ML

We first examined responses of face cells to pairs consisting of a face and a non-face object, selected from three different faces and three different objects (Supplementary Figure 2). The two stimuli were presented either horizontally or vertically aligned, each 3.2° away from the fixation point; both possible locations of the face and object were tested (Figure 1a).

**Figure 1.**
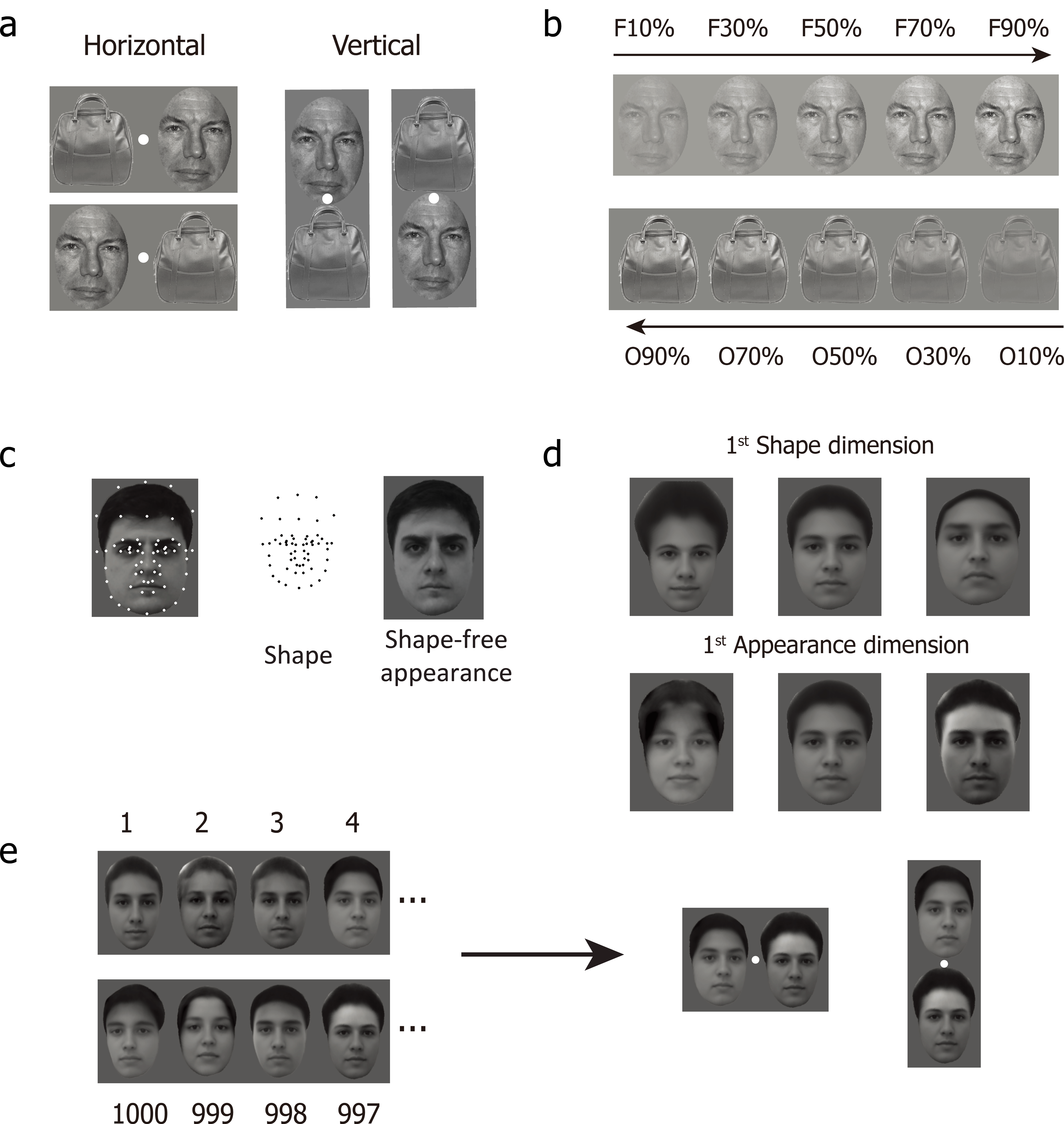
Stimuli for the experiments. (**a**) Two spatial configurations were used in the face-object experiments: horizontal (left) and vertical (right). (**b**) For each configuration and each face-object pair, five different relative contrast levels were tested. (**c**) 58 landmark points were labeled on 200 facial images from a face database (FEI face database; example image shown on left). The positions of these landmarks carry shape information about each facial image (middle). The landmarks were smoothly morphed to match the average positions of the 200 faces, generating an image carrying normalized appearance information about each face (right). (**d**) Facial images corresponding to the first PC for shape (top) and the first PC for normalized appearance (bottom). (**e**) 1000 images were randomly drawn from this space and then paired; four pairs are shown (left). Face pairs were presented in two spatial configurations, horizontal and vertical (right).

For each face-object pair, five relative contrasts of the face and object were tested, from a low contrast face and high contrast object, to a high contrast face and low contrast object (Figure 1b). In area V1, it has been reported that the integration rule can change from averaging to winner-take-all depending on the contrast of the two stimuli presented^8^; we were interested in whether this also holds true in IT cortex. In addition to the five face-object pairs of varying contrast, we also measured responses to the same stimuli presented in isolation, for a total of 180 pair stimuli (4 spatial configurations × 9 face-object identity pairs × 5 contrasts) and 120 isolated stimuli (4 positions × 6 face/object identities × 5 contrasts). Stimuli were presented for 250 ms (ON period) interleaved with a gray screen for 150 ms (OFF period). The same set of 300 stimuli were presented to each cell from 8 to 10 times each. Responses to the stimuli were calculated as the firing rate in a time window 60-220 ms after stimulus onset.

We found that when a face was presented in the contralateral visual field and a non-face object in the ipsilateral field, cells followed a winner-take-all rule: the response to the face-object pairs was very similar to the response to the isolated constituent face, independent of relative contrast (Figure 2a shows an example cell, Figure 2b shows the population average). To quantify the integration rule, we assumed that cells are performing weighted averaging: *R_pair_* = *wR_face_* + (1 − *w*) *R_object_* and we computed w, the weight of the face response, for each cell. For a cell following an averaging rule, *w* = 0.5; for a cell following a winner-take-all rule that responds more to faces, *w* = 1. When a face was present in the contralateral visual field and a non-face object in the ipsilateral visual field, w was close to 1 for all contrasts (Figure 2c).

**Figure 2.**
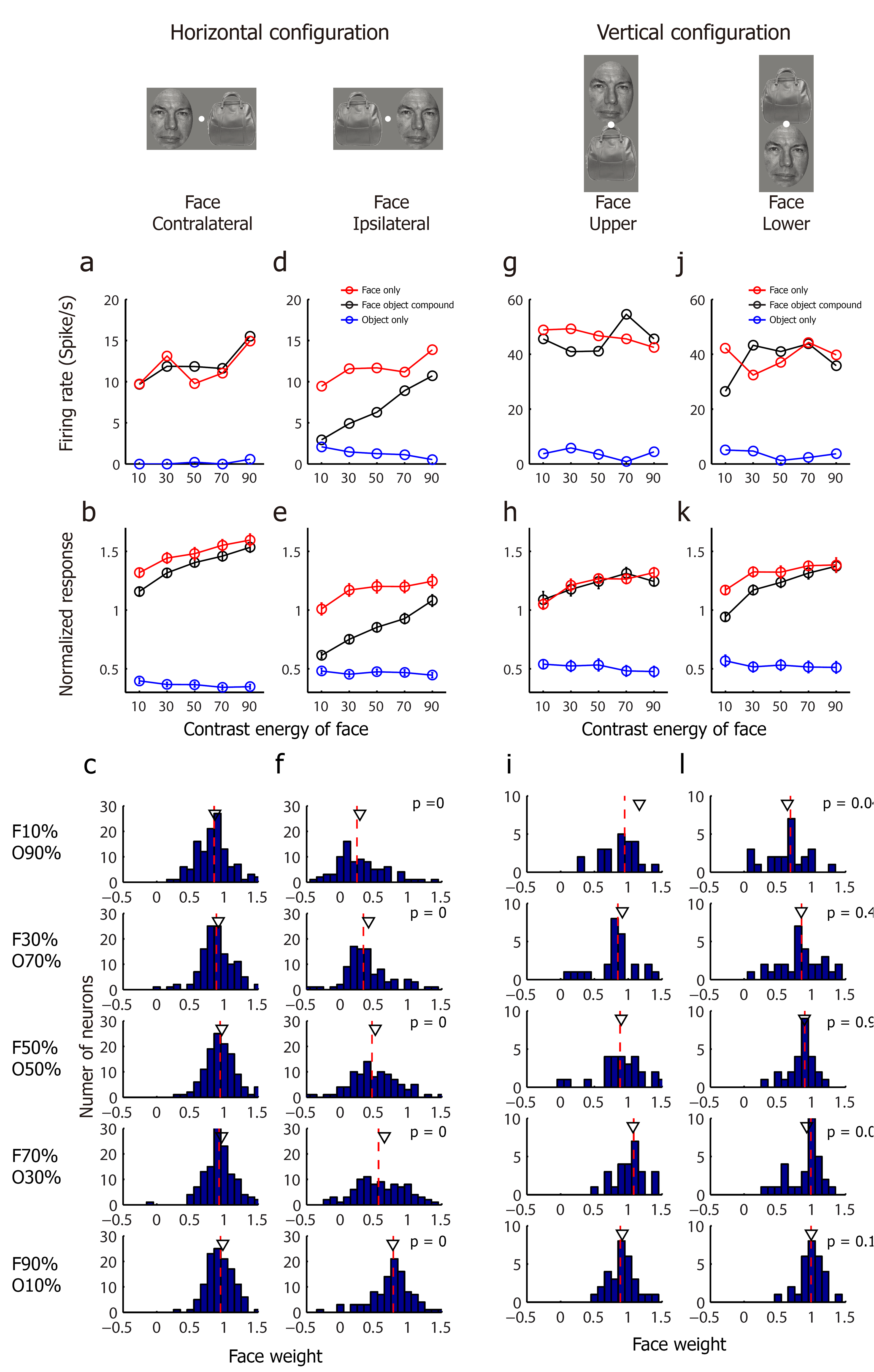
ML responses to face-object pairs. (**a**) Responses of one example neuron to face-object pairs with face presented in the contralateral visual field and object presented in the ipsilateral visual field. Black lines represent the responses to the face-object pairs with at five different relative contrast levels. Red (blue) lines represent the responses to the constituent face (object) of the corresponding face-object pair when presented in isolation. (**b**) The population mean response to the same condition as in (a). For each cell, responses were normalized by the mean response to all stimuli. Error bars denote the S.E. across different cells. (**c**) Population distributions of the face weight (see Methods) computed at five relative contrast levels. The red dashed line denotes the median value of the distribution, which are 0.84, 0.88, 0.96, 0.93, 0.95 (from top to bottom). The black triangle denotes the mean value of the distribution. (**d-f**) Same as (a-c), for the condition in which faces were presented in the ipsilateral visual field and objects were presented in the contralateral visual field. The median values in b3 are 0.21, 0.34, 0.46, 0.61, 0.78. P values displayed in (f) indicate the t-test significance value comparing the population distributions of the face weights between the two horizontal conditions for each contrast level. (**g-i**) Same as for (a-c), for the condition in which faces were presented above the fixation point and objects were presented below the fixation point. The median values in c3 are 0.96, 0.85, 0.89, 1.08, 0.89. (**j-l**) Same as for (a-c), for the condition in which faces were presented below the fixation point and objects above the fixation point. The median values in d3 are 0.68, 0.85, 0.90, 0.98, 0.99. P values displayed in (l) indicate the t-test significance value for comparing the population distributions of the face weights between the two vertical conditions for each contrast level

When a face was presented in the ipsilateral visual field and a non-face object in the contralateral visual field, a very different integration behavior emerged. Now, the response to the face-object pair depended strongly on the relative contrast between the face and the object (Figure 2d, 2e). The weight of the face response increased from around 0 to around 1 as the contrast of the face increased, exactly like what has been found with paired sinewave grating (plaid) experiments in V1^8^. Overall, the results so far show that in response to a face-object pair, the integration behavior used by ML cells is highly dependent on the spatial arrangement and relative contrast of the constituent face and object. The same cell can switch between winner-take-all and weighted averaging (Figure 2a-f; data for three monkeys shown separately in Supplementary Figure 3a).

**Figure 3.**
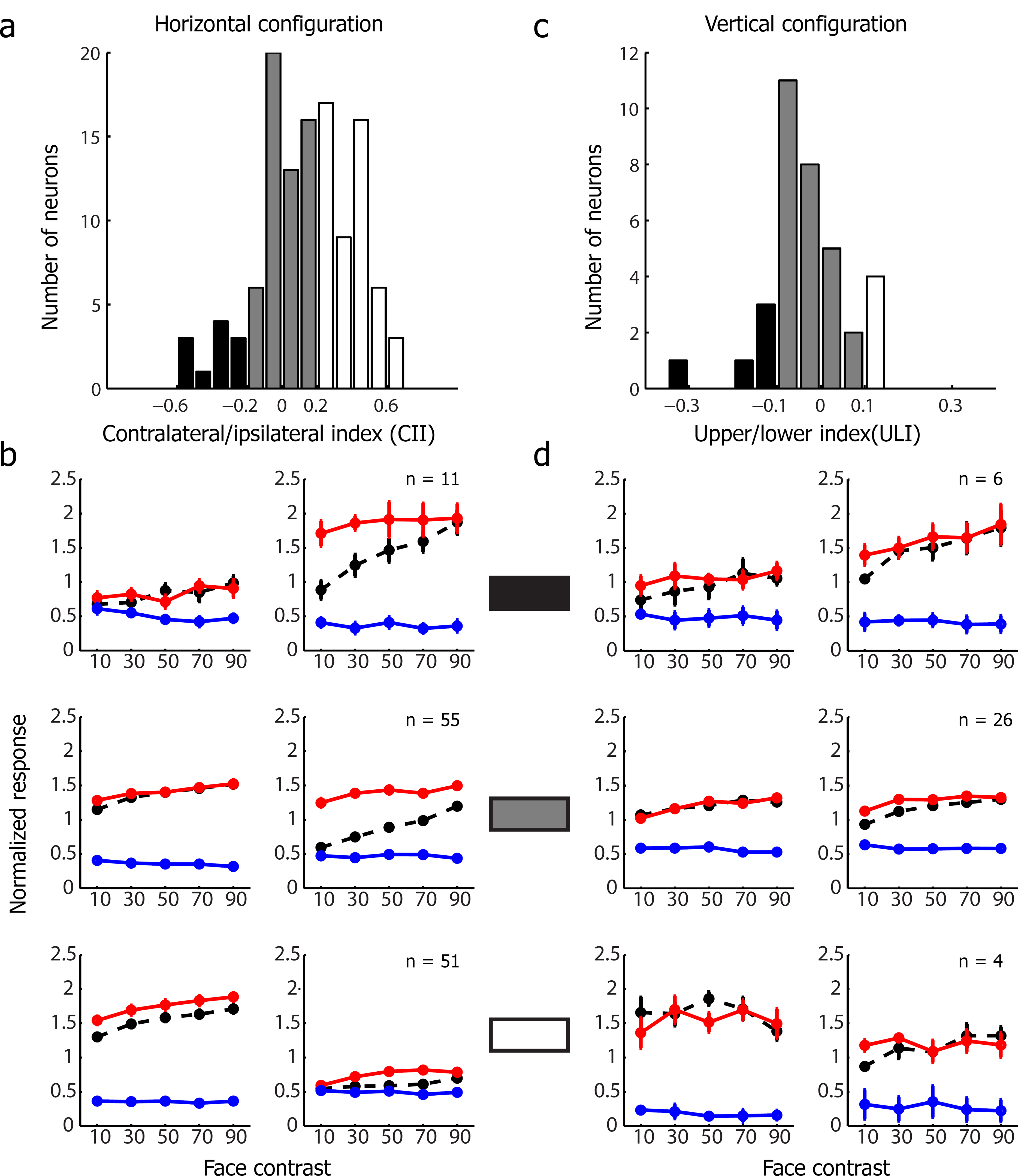
The integration rule for face-object pairs does not depend on a neuron’s spatial tuning. (**a**) The distribution of preference for contralateral/ipsilateral across the population, defined as CII (see Methods). Neurons were classified into three groups: (1) high preference for the ipsilateral visual field (CII < −0.2, black), (2) low preference or no preference (|CII| ≤ 0.2, gray), (3) high preference for the contralateral visual field (CII>0.2, white). (**b**) The population mean response of each group defined in (a). Conventions as in Figure 2a. (**c**) The distribution of preference for upper/lower visual field across the population, defined as ULI (see Methods). Neurons were classified into three groups: (1) high preference for the upper visual field (ULI < −0.1, black), (2) low preference or no preference (|ULI| ≤ 0.1, gray), (3) high preference for the lower visual field (ULI>0.1, white). (**d**) The population mean response of each group defined in (c). Conventions as in Figure 2a.

Could the finding that the response to a face-object pair always followed a winner-take-all rule in the hemisphere contralateral to the face be a consequence of the spatial tuning of ML cells? For example, if neurons in ML have exclusively contralateral receptive fields, then it might not be surprising for them to follow a winner-take-all rule when a face is presented contralaterally (though the weighted averaging rule in the ipsilateral case would still need to be explained).

Receptive field sizes in IT cortex are generally relatively large compared with those in early visual areas V1-V4^15, 27, 28^, and most receptive fields extend across the vertical meridian into both visual hemifields^29^. To clarify, for the specific population of ML neurons we recorded from, how multiple object integration behavior is related to receptive field location, we performed the following analysis: We defined each unit’s preference for contralateral versus ipsilateral isolated faces (contra-ipsi index) as follows: *CII* = (*R_contra face_* − *R_ipsi face_*) / (*R_contra face_* + *R_ipsi face_*). As expected, the population showed a preference for the contralateral visual field (in Figure 3a, the distribution of CII values is significantly skewed to the right (the mean value CII = 0.13 > 0, t-test, p < 10^−4^). Nevertheless, we observed a subpopulation of neurons which showed a strong preference for the ipsilateral visual field. We divided the whole population into three groups: cells with high preference for the contralateral visual field (CII > 0.2), cells with no/low preference (|CII| ≤ 0.2), and cells with high preference for the ipsilateral visual field (CII < −0.2). We then analyzed the integration rule for each of these three groups separately (Figure 3b). All three groups showed similar integration behavior: units performed winner-take-all when a face was in the contralateral visual field and weighted averaging when a face was in the ipsilateral visual field. This suggests that the multiple object integration behavior observed in ML does not depend on a particular neuron’s spatial tuning, but is a general property of ML.

Another way to address the influence of receptive field location on integration behavior is to present the two stimuli aligned vertically instead of horizontally: most cells in ML respond equally well to faces above and below fixation. Furthermore, this would allow direct comparison to a previous study^4^ reporting that most IT cells follow a simple averaging rule (with equal weights for both stimuli), which used vertically aligned stimuli. Thus we next analyzed responses to face-object pairs aligned vertically around the fixation point (Figure 1a, right). We found that in this configuration, cells followed a winner-take-all rule, regardless whether the face was above or below the fixation point, and regardless of the relative contrast between the face and object (Figure 2g, j show individual examples; Figure 2h, k show population averages; Figure 2i, l show histograms of face response weights). We further confirmed that this behavior did not depend on spatial tuning of neurons for the upper versus lower visual field (Figure 3c, d).

We also tested multiple object integration behavio as a function of the face selectivity of particular neurons. A previous study suggested that IT neurons with high object selectivity should have low tolerance to clutter^13^. Supplementary Figure 4 shows that integration behavior of face cells did not depend on their face selectivity.

Finally, we also computed face weights as a function of time under both spatial configurations, using a 5 ms sliding window. This did not reveal any significant change in integration rule over time (p>0.05, Bonferroni corrected, for all tested time points (0 ∼ 400ms); also see Supplementary Figure 5).

### Response to two preferred stimuli in ML

In the previous experiment, we examined the integration behavior of ML cells for a preferred stimulus (face) paired with a non-preferred stimulus (non-face object). Do ML cells show the same behavior when two preferred stimuli, i.e., a pair of faces, are presented? To address this, we presented 1000 face pairs aligned either horizontally or vertically. We decided to present such a large set of faces in order to cover the full dynamic range of ML cell responses: if we had chosen just three faces, and all three happened to be effective stimuli for a cell, then it would have been impossible to distinguish between averaging and winner-take-all behavior using the responses to these three stimuli.

We selected the 1000 face pairs using a strategy motivated by a recent study from our lab which found that ML cells are strongly tuned to specific dimensions in a realistic face space^30^. Here, we adapted our previous approach of generating realistic face stimuli using an “active appearance model”^31^ as follows: for each of 200 frontal faces from an online face database (FEI face database), a set of landmarks were labeled by hand (Figure 1c, left). The positions of these points carry information about the shape of the face and the shape/position of internal features (Figure 1c, middle). Then the landmarks were smoothly morphed to a standard template (average shape of landmarks; Figure 1c, right); the resulting image carries normalized appearance information. In this way, we extracted a set of 200 shape descriptors and 200 appearance descriptors. To construct a realistic face space, we performed principal components analysis on the shape and appearance descriptors separately, to extract the feature dimensions that accounted for the largest variability in the database, retaining the first three principal components (PCs) for shape and first three PCs for appearance (Figure 1d). This results in a 6-dimensional (6-d) face space, where every point represents a face, obtained by starting with the average face, first adding the appearance transform, and then applying the shape transform to the landmarks. The advantage of generating faces defined by these six dimensions is that it allows us to systematically and evenly explore the entire face space.

To generate stimuli for our experiment, we randomly drew 1000 faces from this 6-d face space. Then we generated 1000 pairs of faces by assigning the i^*th*^ face to the (1001-i)^*th*^ (i = 1,2,3,…1000) face as a pair (Figure 1e). This ensured that all 1000 faces were presented at both positions. In separate experiments, the pairs were aligned either horizontally or vertically around the fixation point, and for each pair, we also measured the responses to the constituent faces presented alone. In this experiment, stimuli were presented for 150 ms (ON period) interleaved with a gray screen for 150 ms (OFF period). The same set of 3000 stimuli for each configuration were presented to each cell from 2 to 4 times each. Responses to the stimuli were calculated as the firing rate in a time window 60-220 ms after stimulus onset.

To quantify neuronal tuning within the 6-d face space, responses of each neuron were first used to calculate a “spike-triggered average” (STA) stimulus^32^, i.e., the average stimulus that triggered the neuron to fire. The STA captures all of the important coding properties of a face cell: by knowing just the STA of a face cell, one can predict almost all of the explainable variance of its response to an arbitrary set of faces^30^. Thus the STA provides a compact characterization of a face cell’s selectivity for faces.

For the horizontal configuration, we calculated the STA for each of the following four conditions: (1) a contralateral face presented in isolation, (2) a contralateral face presented as part of a pair, (3) an ipsilateral face presented in isolation, and (4) an ipsilateral face presented as part of a pair. Figure 4a shows the STAs for these four conditions for four different example cells. The STA shape was very similar for conditions (1) and (2), showing that tuning to a contralateral face doesn’t depend on whether another face is presented. Very surprisingly, for condition (4), we observed almost no tuning, *as if cells became completely blind to the ipsilateral face when a contralateral face was present*, i.e., cells follow a contralateral-take-all rule. Importantly, this was not due to cells having exclusively contralateral receptive fields: cells showed clear tuning to ipsilateral faces presented in isolation (Figure 4a, column three). These results were consistent across the population (Figure 4b; data for three monkeys shown separately in Supplementary Figure 3).

**Figure 4.**
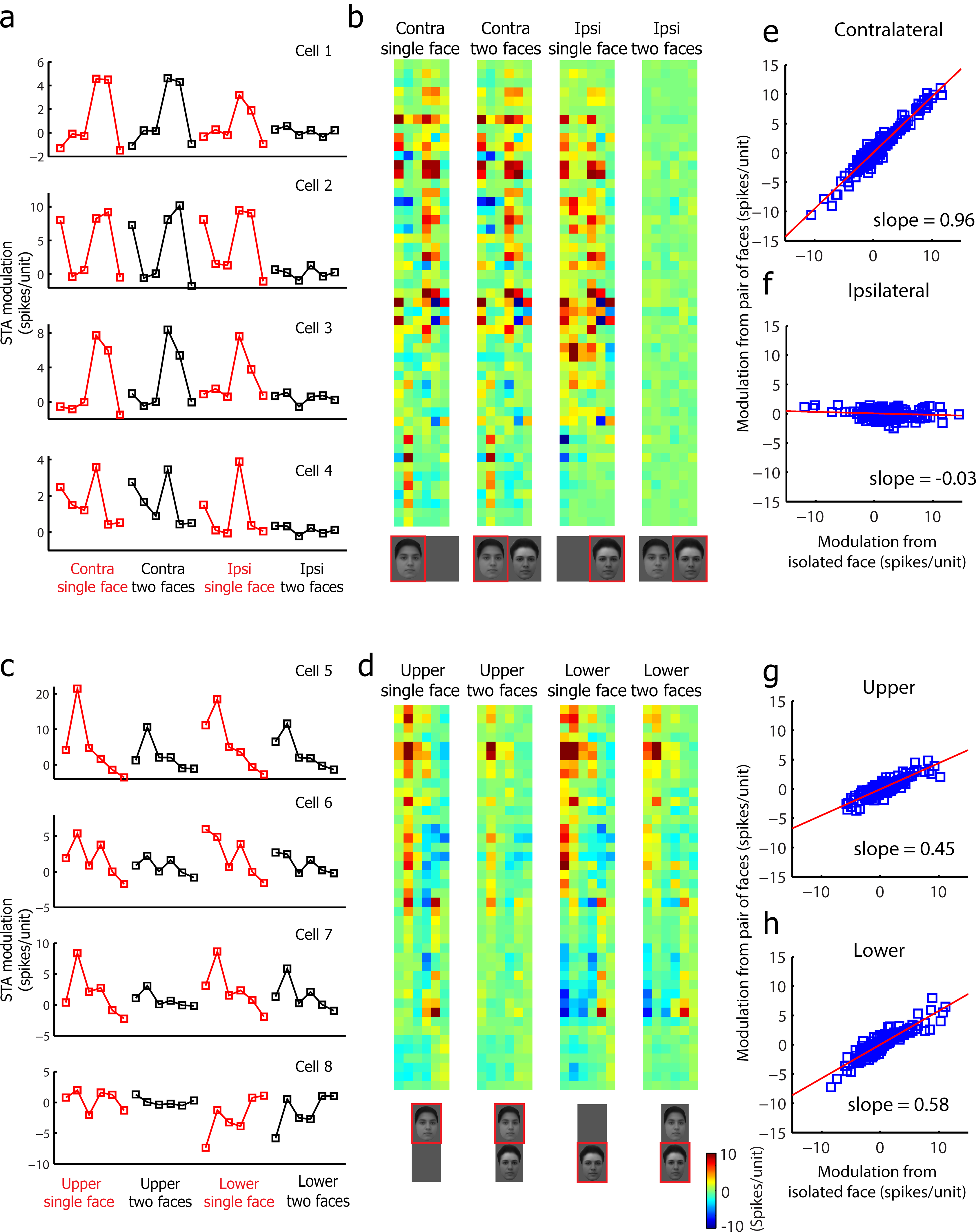
ML responses to face-face pairs. (**a**) STAs for four example neurons, for faces presented in the horizontal configuration. For each neuron, the STA was computed for four different conditions: (1) a contralateral face presented in isolation, (2) a contralateral face presented as part of a pair, (3) an ipsilateral face presented in isolation, and (4) an ipsilateral face presented as part of a pair. (**b**) STAs for all recorded neurons when two faces presented in the horizontal configuration. Each row represents one neuron, and each column represents one face space dimension. (**c-d**) Same as (a) and (b), for the condition in which two faces were presented in the vertical configuration. For each neuron, the STA was computed for four different conditions: (1) an upper face presented in isolation, (2) an upper face presented as part of a pair, (3) a lower face presented in isolation, and (4) a lower face presented as part of a pair. (**e**) STA values from the contralateral face obtained in the single face condition (horizontal) plotted against STA values from the contralateral face obtained in the paired face condition (vertical). The red line indicates the best linear fit of the data. (**f**) Same as (e), for STA values from ipsilateral faces. (**g**) Same as e, for STA values from faces presented above the fixation point. (**h**) Same as (e), for STA values from faces presented below the fixation point.

When two faces were presented vertically, we saw very similar tuning across all four conditions (Figure 4c, d). However, the gain of tuning was smaller for the two paired conditions compared to the two isolated conditions.

To further clarify the correlation between STAs obtained across the four conditions, we plotted the STA values measured in the isolated and paired conditions. Figure 4e shows a scatter plot of STA values measured for contralateral faces presented in a pair versus contralateral faces presented in isolation; each cell contributes six points to the plot, corresponding to the six dimensions of the STA. The slope of the plot is 0.96, indicating almost identical STA gain for the two conditions. This suggests that cells are using an exact contralateral-take-all rule in this situation, and not some other rank-preserving interaction for generating clutter invariance ^9^. Figure 4f shows a scatter plot of STA values measured for ipsilateral faces presented in a pair versus ipsilateral faces presented in isolation. The slope is −0.03. Figure 4g, h shows scatter plots of STA values measured for above/below-fixation faces presented in a pair versus above/below-fixation faces presented in isolation. The slope of the two plots are 0.45 and 0.58, close to the value of 0.5 expected for cells following an averaging rule. Overall, the experiments with two faces show that cells switch between a contralateral-take-all rule and an averaging rule, depending on whether the faces are aligned horizontally or vertically.

So far, for the two face experiment, we have examined how tuning characterized by the STA changes when two faces are presented compared to when a single face is present. We also analyzed absolute response magnitudes to the 2000 face stimuli across the different conditions (Supplementary Figure 6). We found that for most cells, the response magnitude to a pair of faces was significantly correlated to the response magnitude to a contralateral face presented alone, but was not correlated to the response magnitude to an ipsilateral face presented alone (Supplementary Figure 6a shows a single cell example, and Supplementary Figure 6c shows population results). For vertically aligned faces, we found that the response magnitude to a pair of faces was significantly correlated to the response magnitude to both upper and lower face presented alone (Supplementary Figure 6b,d).

### A parsimonious explanation for integration behavior

So far, our results suggest that single ML cells switch between a diverse set of behaviors for multiple object integration: for a particular cell in ML, responses to pairs of objects can be described by winner-take-all, contralateral-take-all, or weighted averaging, with the invoked behavior depending on the category, spatial configuration, and relative contrast of the two objects.

At first glance, this may seem magical. How can a cell infer the particular visual context in order to select the appropriate behavior? Is there a unified explanation for these diverse integration behaviors? Below, we show how all of the results can be explained by the canonical neural computation of normalization, which has been observed in many different systems (vision, olfaction, audition) across multiple species ^25^. Normalization refers to an operation in which the responses of neurons are divided by a common factor representing the summed activity of a pool of neighboring neurons. We show that to explain the present results regarding multiple object representation in IT within the normalization framework, the only ingredient that needs to be added is the homogenous category selectivity of neighboring neurons forming the normalization pool. In our normalization model, we assume that the response of a cell to two objects is given by the following formula:

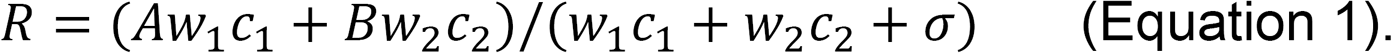

Here, *R* is the response of the cell to a pair of objects with contrasts *c*_1_ and *c*_2_, *A* is the response to object 1 alone at high contrast (i.e., *c*_2_ = 0, *c*_1_ ≫ *σ*), *B* is the response to object 2 alone at high contrast, *w*_1_ represents the weighting of neighboring neurons (i.e., normalization pool) for objec 1, and *w*_2_ represents the weighting of neighboring neurons for object 2.This equation is identical to Equation 9 in Carandini and Heeger (2013), with one new ingredient: the weighting terms *w*_1_ and *w*_2_ (note: we use “A” and “B” to represent responses to objects 1 and 2, instead of “*w*_1_” and *w*_2_” as in Carandini and Heeger (2013), since we use “*w*_1_” and “*w*_2_” to represent normalization weights). The weighting terms *w*_1_ and *w*_2_ endow normalization with an extra degree of freedom, such that the strength of normalization can vary depending on the category and spatial location of the two objects. The justification for this is that we are assuming the normalization pool is not only determined by the contrast of the two stimuli being integrated but also by the category and spatial selectivity of the neighboring neurons. For example, a cell in a face patch should experience more suppression by a face than by a non-face object, even if they have the same contrast, because there are more cells selective for faces than non-face objects in the normalization pool; in our normalization equation, this would be expressed by *w_face_* ≫ *w_object_*.

In Figure 5a-c and Table 1, we show how simple, reasonable assumptions about the normalization factors associated with contralateral faces (*w*_1_), ipsilateral faces (*w*_2_), contralateral objects (*w*_3_), and ipsilateral objects (*w*_4_), namely, *w*_1_ ≫ *w*_2_ ≈ *w*_3_ ≫ *w*_4_, can explain all of the horizontal configuration results, and a similar set of assumptions can explain the vertical configuration results as well. Importantly, these assumptions are experimentally supported by measurements of LFP response magnitudes from ML to the four different conditions, for both spatial configurations (Figure 5d, e). The LFP is thought to measure synaptic activity in thousands of neurons near the electrode tip^33^, and therefore provides a reasonable estimate of the pooled suppressive inputs for each of the four conditions. We used LFP amplitudes at the highest contrast as a proxy for the weights in the normalization model, and found that responses based on the model fits were highly correlated to the actual responses: *r* = 0.994 (p < 10^−4^) for horizontal configuration and *r* = 0.987 ( p < 10^−4^) for vertical configuration (Supplementary Figure 7). As a sanity check, in Figure 5f, g we fit our face-object data to the normalization model to obtain quantitative estimates for the values of *w*_1_ - *w*_4_. These values agreed well with the approximations we obtained from our LFP measurements (compare Figure 5d, e with Figure 5h, j). We also used LFP amplitudes at different contrasts as proxy for the product weight * contrast in the normalization model (Figure 6a, b), and found that responses based on the model fits were highly correlated to the actual responses (Figure 6c-f). Overall, these results show how the widely-accepted framework of normalization^25^ can be extended to explain multiple object representation in IT cortex, with one added assumption that homogenous category selectivity of neighboring neurons forming the normalization pool produces different normalization weights for faces compared to objects within a face patch.

**Table 1.** Summary of multiple object integration rules in ML across different stimulus conditions. Each of the observed rules follows directly from the normalization model presented in Figure 5. For example, for a contralateral face (*w*_1_) and an ipsilateral object (*w*_2_) presented in the horizontal configuration, since w _1_is much larger than *w*_2_ (Figure 5C), we deduce that R follows a winner-take-all rule (Figure 5b, second line).

| Horizontal Configuration |  |  | Vertical Configuration |  |  |
| --- | --- | --- | --- | --- | --- |
| Contralateral | Ipsilateral | Rule | Upper | Lower | Rule |
| Face | Object | Winner-take-all | Face | Object | Winner-take-all |
| Object | Face | Weighted-average | Object | Face | Winner-take-all |
| Face | Face | Contralateral-take-all | Face | Face | Average |

**Figure 5.**
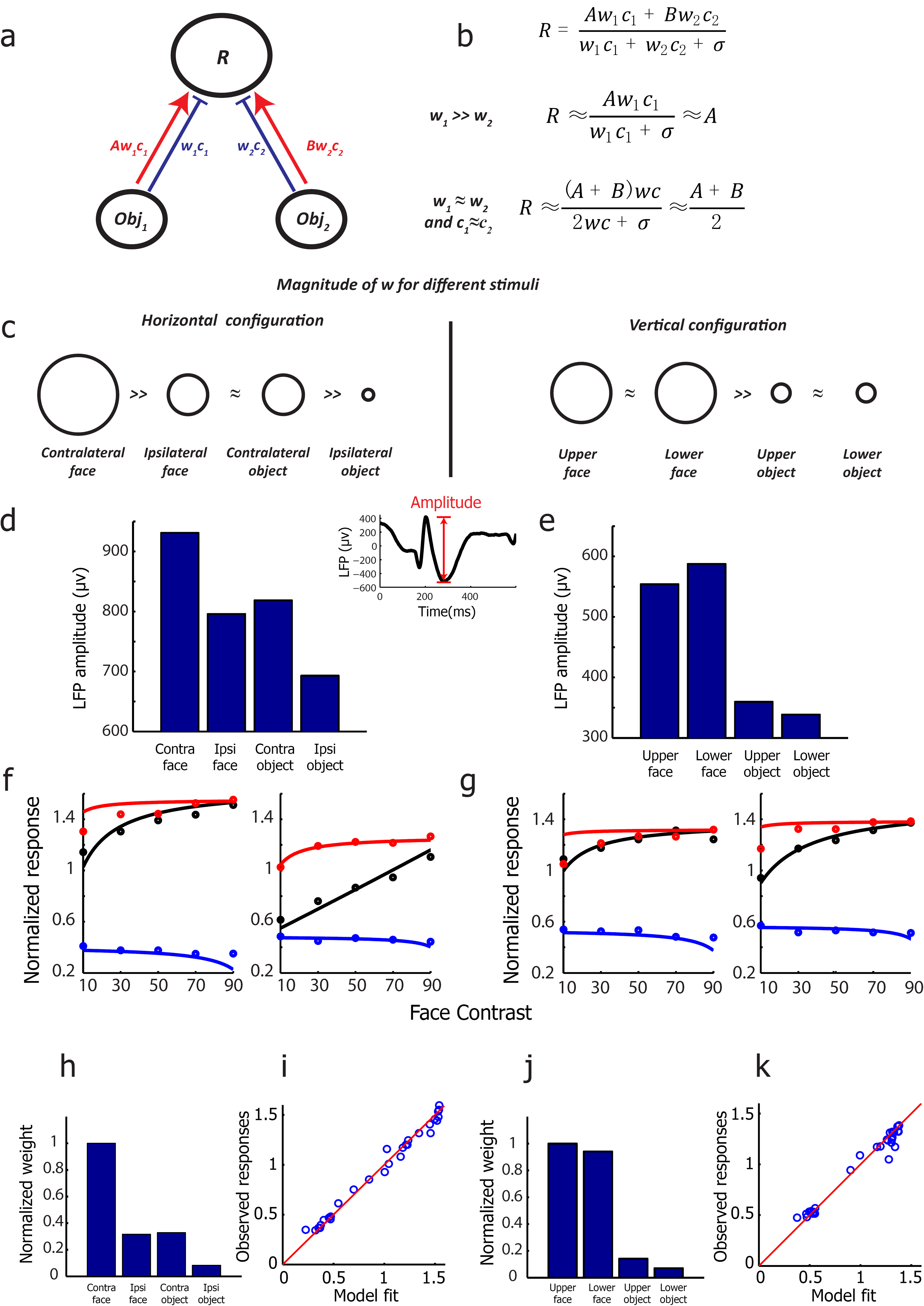
A normalization model can explain diverse ML integration rules. (**a**) Schematic normalization model for an IT cell responding to two objects (adapted from Reynolds et al., ^11^). The red line indicates the excitatory input, and the blue indicates the inhibitory input. R indicates the firing rate of the cell; *w*_1_, *w*_2_ indicate the strength of the inhibitory input associated with the two objects; c_1_, c_2_ indicate the contrast energy of the two objects; A, B represent the ratios between the excitatory and inhibitory input for the two objects. (**b**) According to the normalization model, the response R can be written as a ratio between the summed excitatory input and the summed inhibitory input. When *w*_1_ is much larger than *w*_2_, the response approximates winner-take-all. When *w*_1_ is similar to *w*_2_ and the contrast of two stimuli are same, the response approximates averaging. (**c**) Estimates of the strength of the inhibitory input for different stimuli (indicated by the size of each circle), based on the strong selectivity for faces versus objects in ML and the spatial tuning properties of ML. Combining these estimates with the normalization equation in (b) allows prediction of the integration rules used by ML cells across all stimulus conditions tested in this paper (Table 1). (**d**) The average LFP amplitude for four conditions (contralateral face, ipsilateral face, contralateral object, ipsilateral object) obtained from the face-object experiment when the face and object were presented in isolation at highest contrast level. The LFP amplitude was defined as the difference between negative peak and positive peak in a time window of 150-350 ms after stimulus onset. **(e)** The average LFP amplitude for four conditions (face in the upper visual field, face in the lower visual field, object in the upper visual field, object in the lower visual field) when the face and object were presented in isolation with highest contrast level. **(f)** The normalization model fit to the face-object experiment when faces and objects were presented horizontally (see Methods for details). Line indicates model fit, circles indicate observed data (same as Figure 2a2, b2). **(g)** Same as (f), for faces and objects presented vertically. **(h)** Relative weights (normalized by the maximum weight) obtained from model fits for faces and objects in (f). **(i)** The correlation between the observed responses and the responses based on the normalization model fit when faces and objects were presented horizontally **(j)** Relative weights (normalized by the maximum weight) obtained from model fits for faces and objects in (g). **(k)** Same as (i), for face and objects presented vertically.

**Figure 6.**
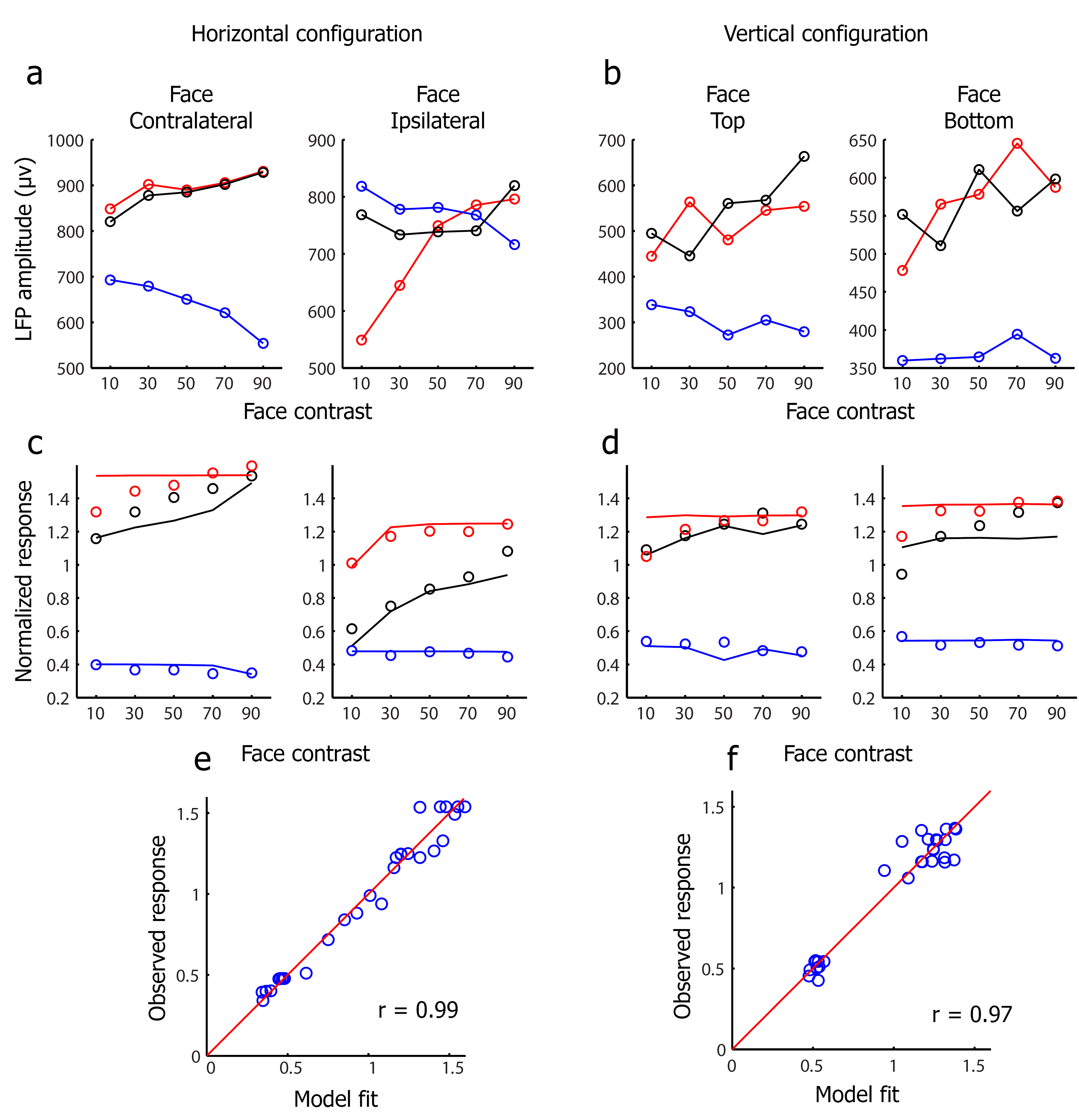
The normalization model using LFP amplitudes across different contrast levels as weights predicts neural responses. **(a)** The LFP amplitudes to face-object pairs when face and object were presented horizontally. Black lines represent the LFP response to the face-object pairs at five different relative contrast levels. Red (blue) lines represent the LFP response to the constituent face (object) of the corresponding face-object pair when presented in isolation. **(b)** Same as (a) to face-object pairs when face and object were presented vertically. **(c)** The normalization model fit using LFP amplitudes across different contrasts as weights to the face-object experiment when faces and objects were presented horizontally (i.e., in the model, instead of using the weight multiplied by the contrast, here we used the amplitudes of the LFP across different contrasts as the weights in the model to fit the data). Line indicates model fit, circles indicate observed data (data same as Figure 2a2, b2). **(d)** Same as (c), for faces and objects presented vertically. **(e)** The correlation between the observed responses and the normalization model fit when faces and objects were presented horizontally. We added a free LFP baseline parameter to obtain the model fits (i.e., the weights in the normalization model were calculated as the LFP amplitude minus the baseline). **(f)** Same as (e), for face and objects presented vertically.

### Integration rules used by cells in the middle body patch MB

Does the normalization model generalize beyond face patches? To test this, we performed recordings in the middle body patch, a region in the lower bank of the superior temporal sulcus (STS) containing a high concentration of body-selective cells (Figure 7a). We presented pairs of bodies and non-body objects (including faces) (Figure 7b). Consistent with previous studies^17^, we found a high concentration of body-selective cells (Figure 7c). In the middle body patch, we found that when a body was presented contralaterally, and a face was presented ipsilaterally, cells showed winner-take-all behavior (Figure 7d). When the body was presented ipsilaterally, and a face was presented contralaterally, cells showed averaging behavior. This is exactly analogous to the face patch, confirming the generality of the normalization model for explaining multiple object integration rules in IT.

**Figure 7.**
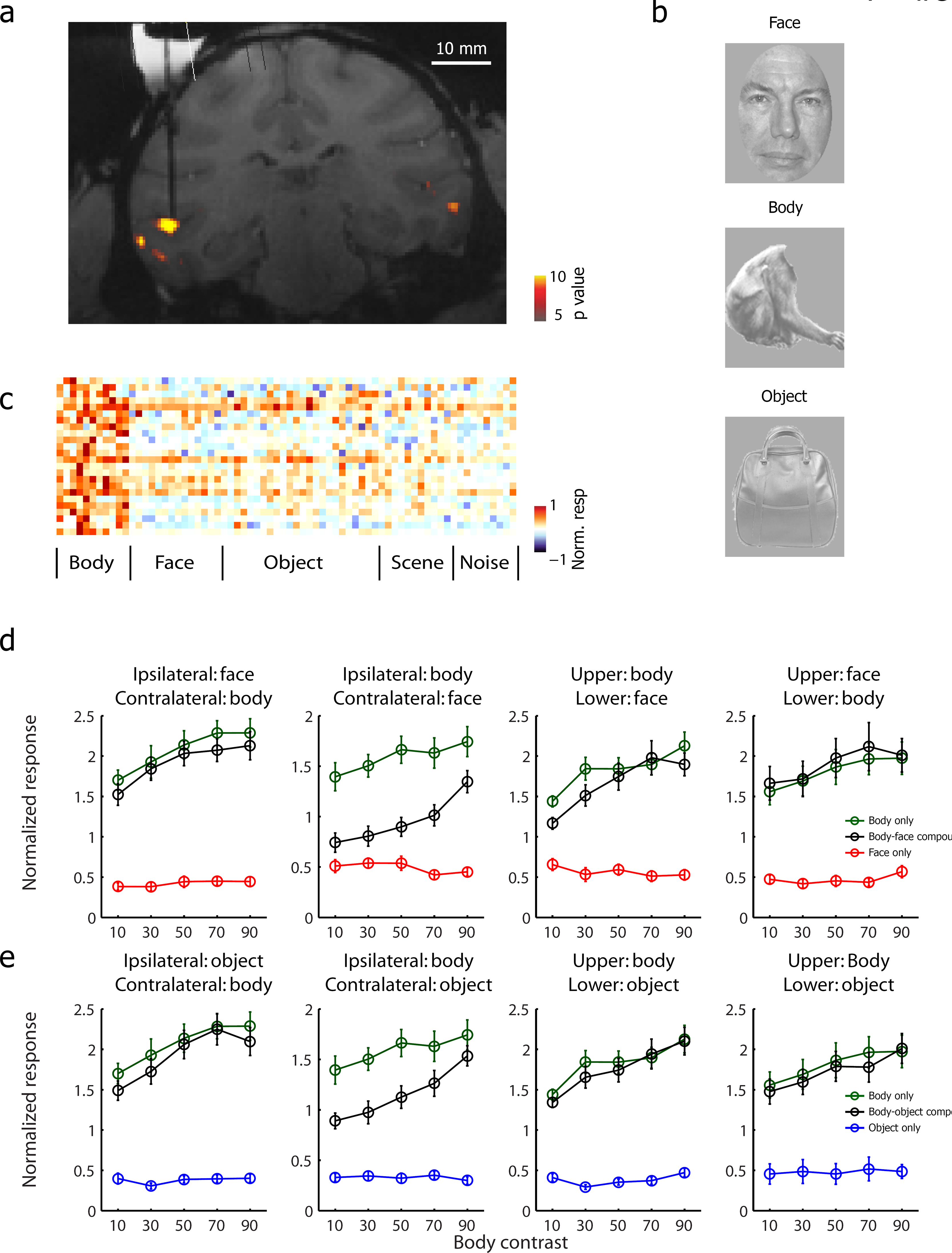
Integration rules in the middle body patch. (**a**) Coronal slice showing the location of the fMRI-identified middle body patch in one monkey targeted for recording; dark black line indicates electrode. (**b**) Example stimuli used to probe integration rules in the middle body patch. **(c)** Mean baseline-subtracted responses of neurons recorded in the middle body patch to stimuli from different object categories. (**d**) Mean response of cells in the middle body patch to a face and a body, presented in the configurations indicated within each panel. Black lines represent the responses to the body-face pairs at five different relative contrast levels. Green (red) lines represent responses to the constituent body (face) of the corresponding body-face pair when presented in isolation. (**e**) Same as (c), for body-object pairs. Analogous to the face patch, body cells followed a winner-take-all rule for bodies in the contralateral visual field, and an averaging rule for bodies in the ipsilateral visual field. Body cells also followed a winner-take-all rule for bodies in the upper/lower visual fields when presented together with a non-body object.

Furthermore, this result also shows that *the integration behavior governing the response to a face and a body is different in a face patch compared to a body patch*. In a face patch, when a non-face object (e.g., a body) is presented contralaterally, and a face ipsilaterally, cells show averaging behavior (Figure 2d, e), not winner-take-all as in the body patch (Figure 7d, leftmost panel). Thus specific integration behaviors depend critically on the specific patch being recorded from (though the general principle of normalization holds across all patches).

One might worry that when a non-face object (e.g., a body) is presented contraterally, and a face ipsilaterally, a face patch cell will generally respond more strongly to the ipsilateral than the contralateral stimulus, whereas a body patch cell will show the reverse pattern, and this might be the source of the different integration behaviors observed in the two patches. To control for this, we identified a small group of face patch neurons (N = 25) which showed a larger response to the contralateral non-face object compared to the ipsilateral face. For these neurons, we still observed an averaging rule when the face was ipsilateral and the object contralateral (Supplementary Figure 8). Thus integration behaviors truly are different in different sub-regions of IT cortex. It is critical to know whether one is recording in a face patch or body patch to understand the integration behavior: it is not sufficient to know only the selectivity of the cell one is recording from.

## Discussion

The effortlessness with which we recognize objects in the cluttered natural world requires explanation. Many studies have explored the role of attention in this process^20, 34^. We tackled the question of how IT cortex integrates responses to multiple objects during passive fixation through targeted recordings in face patch ML and body patch MB. Contrary to previous studies^4, 6^, we found clear evidence for winner-take-all behavior in both of these category-selective regions. It is intuitively obvious that winner-take-all behavior for multiple object integration should aid clutter-invariant recognition. In the section “*Benefits of normalization in a homogeneous patch*” of the Methods, we confirm this through explicit computational modeling, showing that object classification performance in clutter using a winner-take-all rule is always better than that for an averaging rule, and the difference is especially large under conditions of low noise and sparse readout (Supplementary Figure 9). Thus our results suggest that category selectivity, by enabling winner-take-all integration under certain conditions through normalization, could play an important role in solving the clutter invariance problem.

Specifically, we found that in face patch ML, when a face and a non-face object were presented simultaneously, in most cases winner-take-all best described the response to the stimulus pair. This was true for faces presented in the contralateral, upper, and lower visual fields. The only exception occurred when a face was presented ipsilaterally and an object contralaterally: in this case, the response to the stimulus pair was best described by weighted averaging, with weight dependent on the relative contrast of the face and object. Our finding of winner-take-all behavior in face patch ML is consistent with previous human fMRI studies exploring multi-object coding in category-selective brain areas^20, 21, 24^.

When two faces were presented simultaneously, the integration behavior in face patch ML depended on whether the faces were presented horizontally or vertically. For the horizontal case, cells followed a contralateral-take-all rule: STA analysis revealed that the response to the face pairs was modulated exclusively by the contralateral face. For the vertical case, cells followed a simple averaging rule, with approximately equal weights of 0.5 for the two faces.

It is important to note that we did not observe winner-take-all behavior in *all* conditions. Thus one might wonder to what extent the physiological findings can really explain clutter-invariant recognition. To explicitly relate the multiple integration rules we observed to human psychophysical studies of recognition in clutter, we quantatively modeled face identification behavior for different stimulus configurations, using the integration rules uncovered in this study. In the section “*A model of face decoding for pairs of faces*” of the Methods, we build a population decoding model to explicitly predict face identity decoding performance when two faces are presented in two configurations (horizontal, vertical). The model shows that when two faces are presented horizontally across the vertical midline, feature values of contralateral faces can be decoded very well, while feature values of ipsilateral faces cannot be decoded at all. When two faces are presented in a vertical configuration, decoding of both faces suffers due to the averaging rule (Supplementary Figure 10).

These behavioral predictions are consistent with several psychophysical findings concerning peception of faces in clutter. First, the results are consistent with a human psychophysical study^35^ investigating the perception of facial expressions of face pairs, which found perceptual averaging of facial expressions when two faces were presented vertically aligned within the same visual hemifield, but no averaging effect when the two faces were presented in opposite hemifields. The computational model of face decoding behavior based on our physiological results is also consistent with the psychophysical observation that the left and right visual hemifields process stimuli separately: in a working memory task, increasing the number of distractor stimuli impedes task performance within each hemifield independently^36, 37^. If there is winner-take-all or contralateral-take-all, then processing of contralateral preferred stimuli becomes impervious to presence of stimuli in the ipsilateral visual field. Finally, our results provide a mechanistic explanation for the phenomenon of face “pop-out”, i.e., the finding that detection of a face is impervious to the presence of distractor objects^38^. In our normalization framework, this is explained by the fact that *w_face_* ≫ *w_object_* within a face patch. Thus overall, we believe that the match between the stimulus integration properties of face cells revealed here and face recognition behavior under various clutter conditions suggests a strong causal link between the former and the latter. However, all our experiments were performed in a passive fixation paradigm. Future work is needed to measure monkeys’ behavior simultaneously with neural responses, to explicitly test whether behavioral clutter sensitivity correlates with that predicted by neural responses.

Results from MB were completely consistent with those from ML. When a body was presented contralaterally, and a face ipsilaterally, cells followed a winner-take-all rule. When a body was presented ipsilaterally, and a face contralaterally, cells followed an averaging rule. While the grand schema was completely consistent between MB and ML, our results also show that *the particular behavior governing the response to a face and a body is different in a face patch compared to a body patch*. In a face patch, when a non-face object (e.g., a body) is presented contralaterally, and a face ipsilaterally, cells follow an averaging rule, not a winner-take-all rule. Thus specific integration behaviors depend critically on the specific patch being recorded from, and it was important for us to know that we were recording in a face versus body patch to make sense of our results. More generally, our findings suggest that for objects besides bodies and faces, it will also be critical to study integration mechanisms in a manner that respects IT functional organization for these objects.

The different integration behaviors exhibited by ML/MB cells can all be explained by the canonical neural computation of normalization, with the added ingredient that normalization of responses to multiple stimuli is weighted by the category and spatial selectivity of neighboring neurons for the stimuli. This simple assumption efficiently captures all of our main findings in both face and body patches (Table 1). Normalization has previously been invoked to explain response adaptation properties of IT cells ^39^. The key conclusion of our paper is that normalization provides *a simple mechanism for cells in a category-selective patch to implement a winner-take-all rule for the preferred object of the patch, and thereby aid clutter-invariant recognition under certain conditions*. In effect, category selectivity provides a “cheap” form of visual attention. From the normalization equation (1), it is simple to see why. A widely-accepted model of visual attention posits that attention acts to change the weights in the normalization equation (1)^40^. For example, *w*_1_ = *w*_2_ = 0.5 would imply equal attention to objects 1 and 2, while *w*_1_ = 1, *w*_2_ = 0, would imply exclusive attention to object 1. In a brain region equipped with a normalization circuit that is category selective for object 1, the cells are *hard-wired* to implement the latter condition. While it would seem to be extremely difficult to implement a mechanism to shut off dendritic inputs representing non-preferred objects at the single-cell level, this behavior arises inevitably in a network of cells with homogenous category selectivity carrying out normalization (Figure 5 and Table 1).

Thus IT cortex is able to readily implement winner-take-all in specific sub-regions. Given how parsimoniously the results in ML and MB could be explained by the normalization model plus the category selectivity of these patches (Figure 5, Table 1), it is plausible that a similar principle governs the response to multiple objects across all of IT. Multiple specialized networks beyond those selective for faces and bodies have been described in IT^41, 42, 43, 44, 45^. By virtue of being spatially clustered, cells in these networks would be expected to implement clutter-invariant integration for their preferred stimulus class under certain conditions.

It is even possible that the need to achieve clutter-invariant recognition could have been an evolutionary driving force for developing category-selective regions. The reason why IT cortex harbors category-selective regions remains unclear. Minimization of wiring length for distinguishing similar objects is frequently offered as one possible explanation^46^. The fact that category-selective regions give rise to a highly desirable computational feature, winner-take-all for the preferred object, suggests an additional possible evolutionary origin for this striking aspect of IT anatomy: enabling clutter-invariant recognition.

A previous study of multiple object representation in IT cortex reported that averaging could explain responses to all stimulus pairs, though a small percentage of cells showed winner-take-all ^13^. A possible reason for the discrepancy is that the non-preferred stimuli used in that study evoked substantial responses in neighboring neurons, leading to suppression of the response to the preferred object, similar to the averaging we observed in our twoface vertical configuration experiment.

If an area is already category-selective, one might wonder why any additional form of filtering is even necessary. The point is that even if the mean population activity within an area is strongly category selective, many individual cells within the area will nevertheless respond significantly to objects from non-preferred categories (e.g., a cell in a face patch detecting faces based on round overall shape might also respond to an apple). Normalization provides a mechanism to filter out these responses in clutter situations.

While the mechanism proposed here for filtering out clutter is less flexible than classic, high-level attention (e.g., we already discussed above how cells in face and body patches show winner-take-all behavior only under certain conditions), it has the advantage of being hard-wired, and hence, constantly in operation. One might wonder how we can be so sure that the integration rules observed in face and body patches are due to bottom-up stimulus-driven rather than top-down attentional effects. After all, it is known that faces can powerfully capture attention in cluttered scenes^47^. We think our data argues strongly for bottom-up stimulus-driven mechanisms for several reasons. First, in the two-face condition, it is unclear how attention can explain the contralateral-take-all rule. The monkey can presumably pay attention to only one face, and that would presumably be the face that wins. But our data shows that the face that wins for a particular cell depends on the hemisphere in which the cell is located. Second, in the face-object condition, it is also unclear how attention can explain winner-take-all in the hemisphere contralateral to the face, but averaging in the hemisphere ipsilateral to the face. If attention were leading to winner-take-all behavior, then we should have also observed winner-take-all in the hemisphere ipsilateral to the face. Third, when we decreased face contrast in the face-object condition, which would be expected to diminish attention to the face, we still saw winner-take-all behavior when the face was contralateral--even at the lowest contrasts. Finally, if the winner-take-all behavior observed in face patch cells could be explained by attention, then in the body patch, we would have expected to see responses to a face-body pair resembling responses to a face presented alone. Instead, we found the exact opposite: when we presented a body and face simultaneously in the vertical configuration, the responses to the face-body pair resembled that to the body presented alone. Together, these arguments show that the integration rules used by face cells and body cells to multiple objects are unlikely to be explained only by top-down attention. Instead, we think the results are most parsimoniously explained by a hard-wired normalization circuit. Of course, it is almost certain that attentional mechanisms act on top of the bottom-up integration rules we observed, to provide additional flexibility in filtering clutter, e.g., in situations where the bottom-up circuit only yields averaging.

Ultimately, we want to understand how we “know what is where by looking,” as David Marr famously defined vision^48^. The responses of IT neurons to multiple objects constitute one important piece of this puzzle, clarifying how the identities of multiple objects are represented, and revealing an important new mechanism by which clutter-invariant recognition can be achieved. Future experiments will need to address how spatial locations of multiple objects are represented, and how the two bodies of information are registered.

## Methods

### Face and Body Patch Localization

All procedures conformed to local and US National Institutes of Health guidelines, including the US National Institutes of Health Guide for Care and Use of Laboratory Animals. All experiments were performed with the approval of the Institutional Animal care and Use Committee (IACUC)_°_

Three male rhesus macaques were trained to maintain fixation on a small spot for juice reward. Monkeys were scanned in a 3T Tim Trio (Siemens, Munich, Germany) while passively viewing images on a screen. Feraheme (AMAG pharmaceuticals) contrast agent was injected to improve signal to noise ratio. Six face selective regions were identified in each hemisphere in both monkeys by identifying regions responding significantly more to faces than to bodies, fruits, gadgets, hands, and scrambled patterns, while three body selective regions were identified by identfitying regions responding significantly more to bodies than to fruits, gadgets, hands and scrambled patterns. Additional details are available in Tsao and Freiwald ^49^, Freiwald and Tsao ^15^ and Ohayon, Freiwald ^50^. In both monkeys, we targeted middle face patch ML located on the lip of the superior temporal sulcus, and the middle body patch MB located on the lower bank of the superior temporal sulcus.

### Single-unit recording

Tungsten electrodes (1–20 Mohm at 1 kHz, FHC) were back loaded into plastic guide tubes. Guide tubes length was set to reach approximately 3–5 mm below the dura surface. The electrode was advanced slowly with a manual advancer (Narishige Scientific Instrument, Tokyo, Japan) and were inserted anew on a daily basis. Neural signals were amplified and extracellular action potentials were isolated using the box method in an on-line spike sorting system (Plexon, Dallas, TX, USA). Spikes were sampled at 40 kHz. All spike data was re-sorted with off-line spike sorting clustering algorithms (Plexon). Only well-isolated units were considered for further analysis. For experiment 1 (face-object pair), we recorded 67 neurons in monkey M1’s right hemisphere, 49 neurons in monkey M2’s right hemisphere and 18 neurons in M3’s left hemisphere. For experiment 2 (face-face pair), we recorded 62 neurons in monkey M1’s right hemisphere, 25 neurons in monkey M2’s right hemisphere, and 6 neurons in monkey M3’s left hemisphere. Results were qualitatively the same across different monkeys and therefore were pooled together for population analyses. For experiment 3 (MB, body-face, body-object pairs), we recorded 14 neurons in monkey M3’s right hemisphere and 8 neurons in monkey M4’s left hemisphere.

### Visual Stimuli and Behavioral task

Monkeys were head fixed and passively viewed the screen in a dark room. Stimuli were presented on a CRT monitor (DELL P1130). Screen size covered 21.6 × 28.8 visual degrees. The fixation spot size was 0.25 degrees in diameter. All images were presented in random order using custom software. Eye position was monitored using an infrared eye tracking system (ISCAN). Juice reward was delivered every 2-4 seconds if fixation was properly maintained. Custom software (Kofiko) was used to present visual stimuli, track fixation, deliver juice, and synchronize stimulus delivery and recording of neural data.

#### Stimuli for face-object experiment

Three different facial identities and three different non-face objects were used for this experiment. The face images are collected under the FERET program. All the raw images were adjusted to have the same mean luminance, same root mean square (RMS) contrast, and same number of pixels; RMS contrast is defined as the standard deviation of the pixel intensities. Two stimulus configurations were used in this experiment (Figure 1a), horizontal and vertical. In the horizontal configuration, a face was placed either contralateral or ipsilateral to the recording hemisphere, while an object was placed on the opposite side. In the vertical configuration, a face was placed above or below fixation, while an object was placed on the opposite side. In both configurations, the center of each image was positioned 3.2 visual degrees from the fixation point. Each object or face spanned 5.6 × 6.4 visual degrees. For each face-object pair, the contrast energy of the face (i.e., the square of the RMS contrast) increased from 10% to 90% in five equal steps, while the contrast energy of the object decreased from 90% to 10% (Figure 1b). As a result, the summed contrast energy of the face-object pair was kept constant across different contrast energy combinations. Stimuli were presented for 250 ms (ON period) interleaved with a gray screen for 150 ms (OFF period). Each stimulus was presented to each cell from 8 to 10 times each.

#### Stimuli for two-face experiments

We used real face images from an online face database, FEI face database (http://fei.edu.br/∼cet/facedatabase.html). This database contains images from 200 individuals. Generation of parameterized face stimuli followed the procedure of previous papers on active appearance model ^31^: First, a set of 58 landmarks were labeled on each of the frontal face images (Figure 1b). The positions of landmarks were normalized for mean and RMS contrast for each of the 200 faces, and an average shape template was calculated. After that, each face was smoothly warped so that the landmarks matched this shape template. This warped image was then normalized for mean and RMS contrast and reshaped to a 1-d vector. Principal component analysis was carried out on positions of landmarks and intensity independently. The first 3 PCs of landmark positions (“shape” dimensions) and the first 3 PCs of intensity (“normalized appearance” dimension) were used to construct a parameterized face space. The distribution of feature values for each PC dimension followed a Gaussian distribution with variance proportional to that of the 200 faces from the database. 1000 images were randomly drawn from this space (and constructed from the 6-d feature vector by inverting the process above). The feature value for each dimension was scaled to have zero mean value and standard deviation of 1. Each face spanned 4.5 × 3.5 visual degrees. The center of each image was positioned 1.75 visual degrees from the fixation point in the horizontal configuration and 2.25 visual degrees from the fixation point in the vertical configuration. Additional details are available in Chang and Tsao^30^.

From these 1000 images, we constructed 1000 pairs for two face images by assigning the i^*th*^ face to the (1001-i)^*th*^ (i = 1,2,3,…1000) face as a pair (Figure 1E). In separate experiments, the pairs were aligned either horizontally or vertically around the fixation point, and for each pair, we also measured the responses to the constituent faces presented alone. Stimuli were presented for 150 ms (ON period) interleaved with a gray screen for 150 ms (OFF period). The same set of 3000 stimuli for each configuration were presented to each cell from 2 to 4 times each.

#### Stimuli for body-object experiment

A body image, a face image, and an object image were used in this experiment. Two stimuli combinations were used in the experiment: 1) body image paried with face image, 2) body image paired with object image. For each combination, the same spatial configurations and timing parameters were tested as in the face-object experiment.

### Data Analysis

#### Face Selectivity Index

The Face Selectivity Index (FSI) (Supplementary Figure 1) was defined by

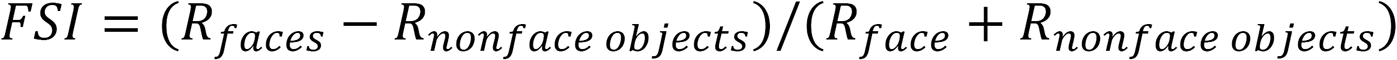

where *R_faces_* is the mean response above baseline to faces and *R_nonfaceobjects_* is the mean response above baseline to non-face objects. An FSI of 0 indicates equal responses to face and non-face objects. An FSI of 0.33 indicated twice as strong response to faces as to non-face objects. For cases where (*R_faces_* > 0) and (*R_nonfaceobjects_* < 0), FSI was set to 1; for cases where (*R_faces_* < 0) and (*R_nonfaceobjects_* > 0), FSI was set to −1.

#### Contralateral ipsilateral index and Upper lower index

##### Contralateral ipsilateral index (CII) was defined as

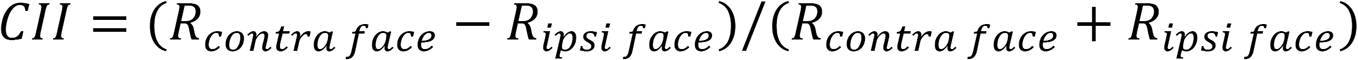

where *R_contra face_* is the neuron’s response to an isolated face presented in the contralateral visual field, and *R_ipsi face_* is the neuron’s response to an isolated face presented in the ipsilateral visual field.

##### Upper lower index (ULI) was defined as

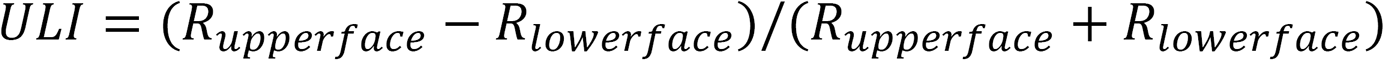

where *R_upper face_* is the neuron’s response to an isolated face presented above the fixation, and *R_lower face_* is the neuron’s response to an isolated face presented below the fixation.

#### Face weight

##### Face weight was defined as

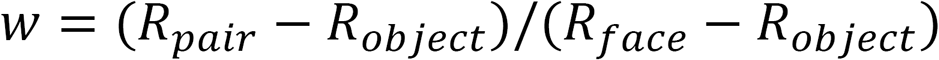

where *R_face_* is the neuron’s response to an isolated face, *R_pair_* is the neuron’s response to a face-pair, and *R_object_* was the neuron’s response to a non-face object.

#### Spike-triggered average analysis

The firing rate in a time window of 60-220 ms after stimulus onset was computed for each stimulus. To estimate the modulation of the each dimenstion, a linear function was fit between the response (i.e., firing rate) and each dimension’s value. The modulation for the dimension was defined as the slope of this linear function. Our definition of spike-triggered average is slightly different from the conventional notion of the average stimulus that triggers a cell to fire: our STAs are proportional to the conventional STA, but give added information about absolute firing rates.

#### Fitting responses to the normalization model (Figure 5)

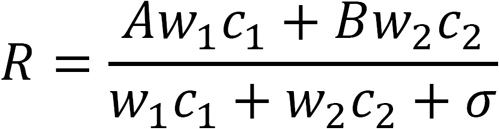

Responses were fit using average data from the face-object experiment (i.e., data points shown Figure 2a2, b2, c2 and d2; Figure 5f,g). For each face-object pair, there are three free parameters, *w*_1_, *w*_2_, and *σ*. *c*_1_ and *c*_2_ are the contrast energies of the face and object. The scaling factors *A* and *B* were set equal to responses to the stimuli presented in isolation at highest contrast level.

### Benefits of normalization in a homogeneous patch

The integration rules used by cells in face patch ML differ markedly from those reported previously based on random recordings in IT^4^. Do the rules uncovered in the present study confer any advantages for object recognition? To address this, following Li et al. (2009), we constructed a hypothetical object space containing three different object categories (A-C) defined along one dimension of object identity (each object was defined by a specific range between −1 and 1) (Figure S10a). We generated a class of hypothetical neurons (N) and simulated the population response to a set of labeled “stimulus scenes” (3000 total: 1000 single object, 1000 two objects, 1000 three objects) following either a winner-take-all or averaging model of multiple object integration. We used these responses to train three linear SVM classifiers to perform object category detection (A/not A, B/not B, C/not C). We then tested these three classifiers on 300 new test images (100 single, 100 two objects, 100 three objects).

The neural response *R* to a stimulus *v* was simulated in the same way as Li et al. (2009):

*R*(*v*) = *H*(*v*) + *c* + *Noise*(*v*), where the neuron’s response function *H* to a single object is *H*(*v*) = *G*(*μ*, *σ*); *μ* is the preferred object of the neuron ( randomly assigned within the stimulus space according to uniform distribution), *σ* specifies the standard deviation of a neuron’s Gaussian tuning (kept at 0.3 in the simulation), and *Noise*(*v*) = *N*(0, *ρ*[*H*(*v*) + *c*]), i.e., response variability is proportional to the response, where *N* is a Gaussian distribution with zero mean and standard deviation proportional to the response with proportionality constant *ρ*. In Li et al., *ρ* was set as a constant (0.25). In our simulation, we tested the simulation results with different *ρ* levels to see how performance varies with different signal/noise ratios.

A second difference between our simulation and Li et al. concerns how we read out object identity. Cox and Riesenhuber (2015) suggest that object recognition may be based on a subpopulation of preferred neurons that respond maximally to the object being recognized and are robust to clutter. Thus, in addition to testing readout of object identity using the full set of neurons (as in Li et al.), we tested a new condition in which readout of object identity used only preferred neurons: to decide whether A is present or not, we only used those neurons which preferred object A to the other 2 objects.

Thus in total, we tested 4 conditions (2 integration rules × 2 readout strategies) at different noise levels *ρ*. We found that performance was always better when applying a winner-take-all compared to averaging rule (Supplementary Figure 10b). The difference was more prominent when the noise level was low. Furthermore, the difference between winner-take-all and averaging rules was larger when sparse readout was applied than when the whole population was read out. These results show that the integration properties of neurons in the categorical-selective patches, arising from normalization in a homogeneous patch, confer a powerful advantage for object recognition in clutter, by making possible a winner-take-all rule. Readout performance can be further enhanced by sparse readout, i.e., reading out only the neurons in the homogeneous patch. Indeed, it is possible that these advantages for oject recognition in clutter may be one of the driving forces for evolution of clustered domains in IT cortex.

### A model of face decoding for pairs of faces

To quantatively model face identification behavior for different stimulus configurations, we built a population decoding model explicitly predicting face identity decoding performance for the scenario in which two faces are presented in two configurations (horizontal, vertical). Our decoding model is based on results from Chang and Tsao (2017), which shows how the identity of a single face can be decoded from responses of a population of face cells. In a nutshell, this earlier work shows that single cells in face patches ML/MF and AM are linearly projecting incoming faces onto their STA. In other words, each cell is performing the computation *r* = *STA* · *F* + *c*, where r is the response of the cell, the *STA* is the 6-d STA vector of the cell, *f* is the incoming face vector, defined by 3 shape and 3 normalized appearance coordinates, and c is a constant offset. If we have a population of face cells, then we can write this equation as: *R* = *T* · *F* + *C*, where *R* is the vector of face cell responses, *T* is the transformation matrix, and *C* is the vector of response offsets. This implies that we can linearly decode *F* by inverting this transformation using responses of a population of face cells: *F* = *T*′ · *R* + *C*′.

To decode the identity of a face presented as part of a pair, we learned the transformation from responses to feature values (i.e., *T*′ and *C*′ above) using the responses of our ML cells to single faces presented ipsilaterally/contralaterally as part of a horizontal pair, or above/below fixation as part of a vertical pair; this resulted in four different (*T*′, *C*′) pairs. We then used these (*T*′, *C*′) pairs to predict the feature values of faces in the corresponding face pair conditions

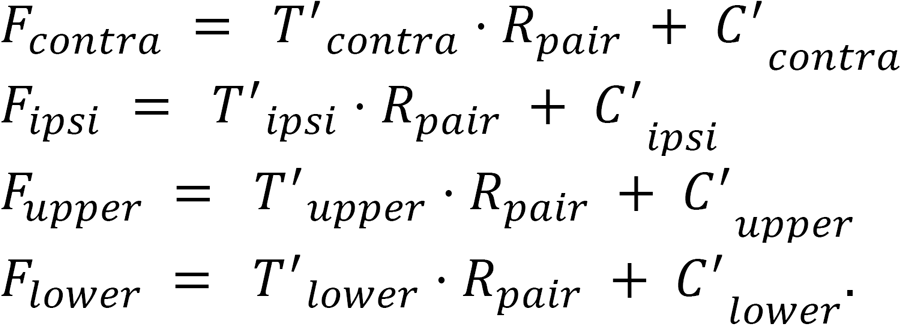

Supplementary Figure 11 shows the results of this decoding model: when two faces were presented horizontally across the vertical midline, feature values of contralateral faces could be decoded very well while feature values of ipsilateral faces couldn’t be decoded at all. When two faces were presented in a vertical configuration, decoding of both faces suffered due to the averaging rule.

## Data availability

The data that support the findings of this study are available from the corresponding authors on request.

---

**Figure S1.**
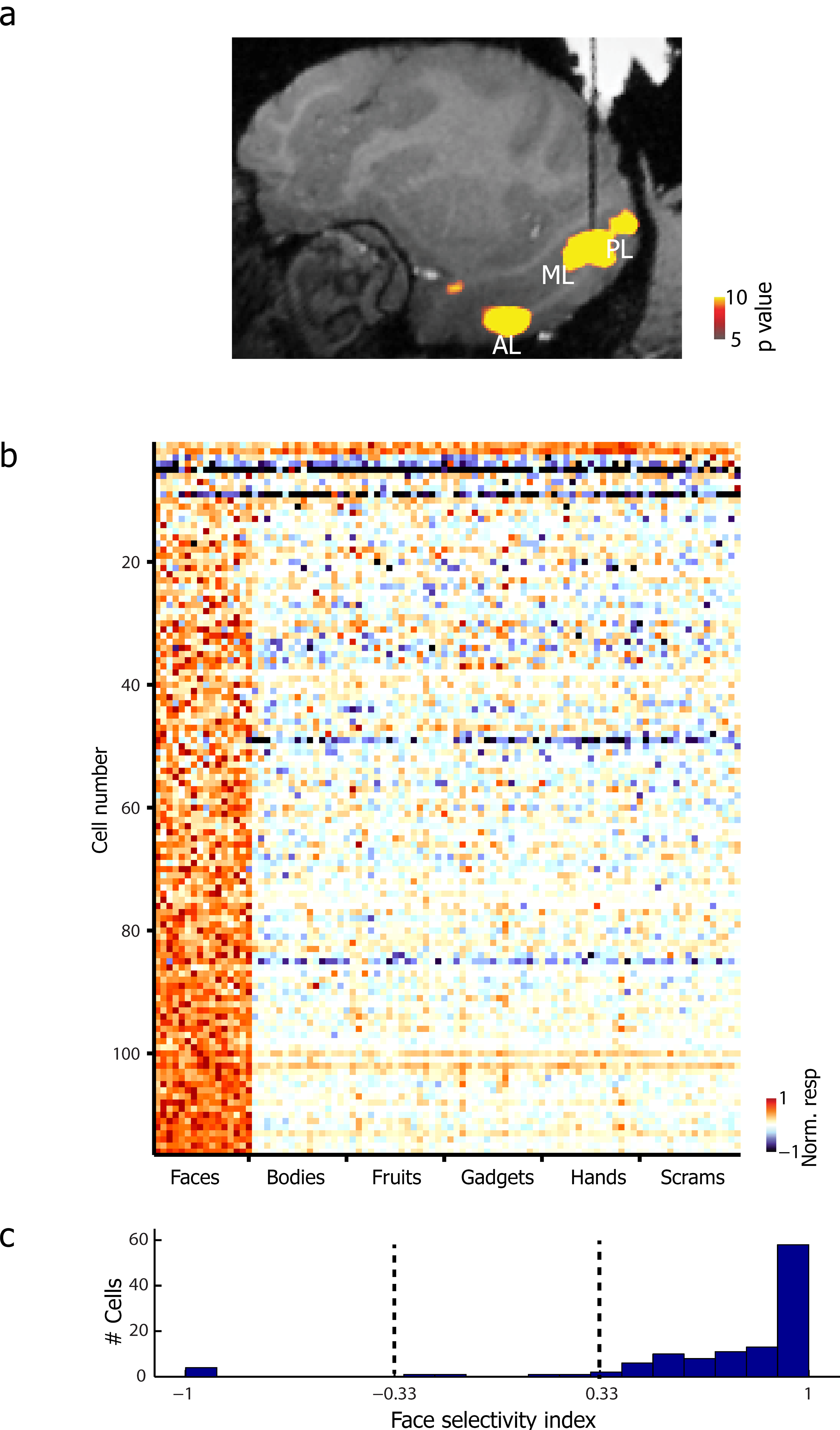
Localization of face patches. (**a**) Sagittal slice showing the location of fMRI-identified face patches in one monkey targeted for recording; dark black line indicates electrode. Stereotactic coordinates for the location of ML were as follows: M1 AP 4.50, ML 25.5 (right hemisphere); M2 AP 4.50, ML 27.43 (right hemisphere), M3 AP 4.00, ML −26.2 (left hemisphere). (**b**) Neuronal responses (baseline-subtracted, averaged from 60 to 220ms) to images of different categories recorded from the ML. (**c**) Distribution of FSI (see Methods) across all visually responsive neurons. Dotted lines indicate |FSI|=0.33.

**Figure S2.**
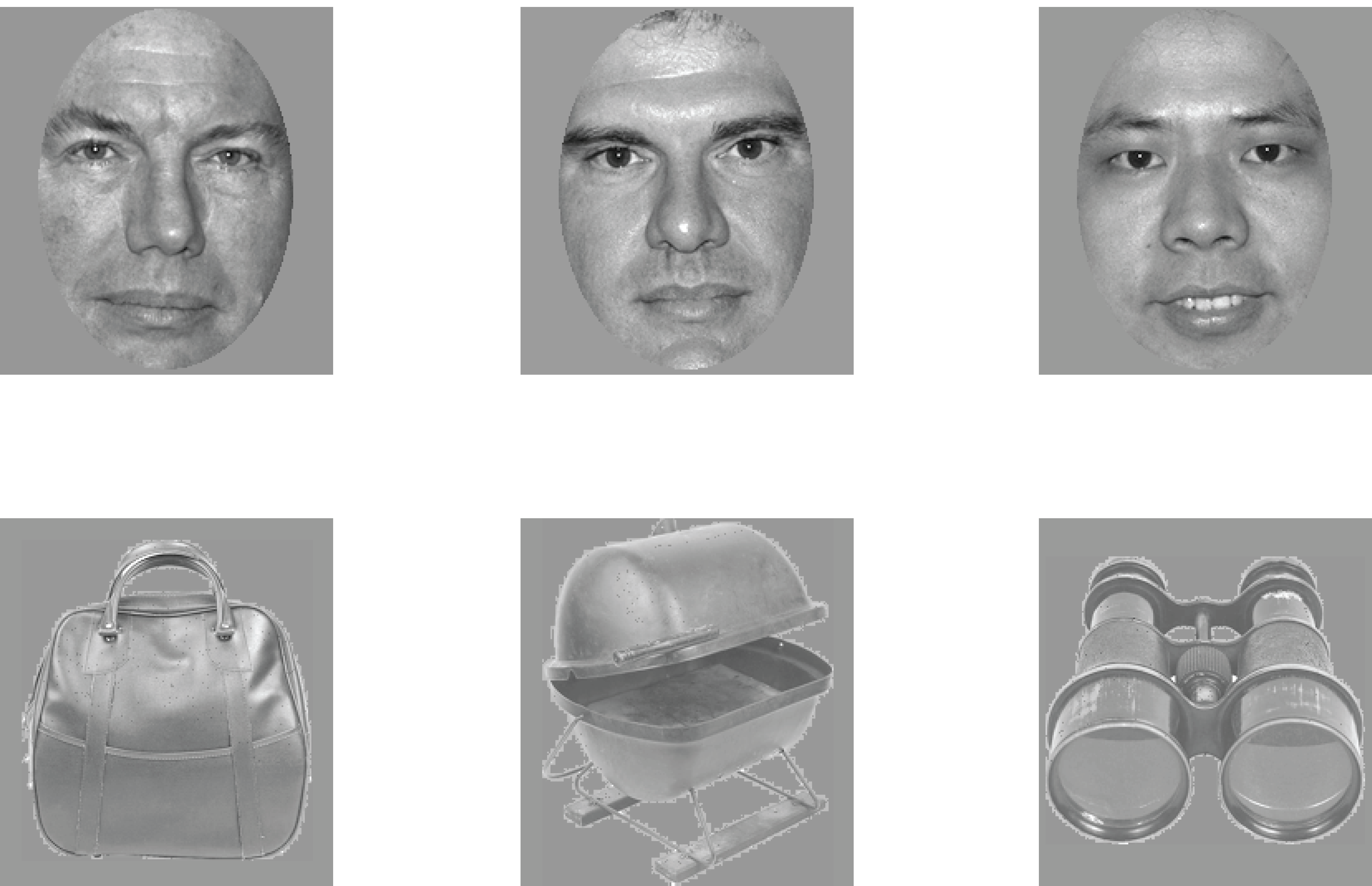
Stimuli for face-object pair experiment.

**Figure S3.**
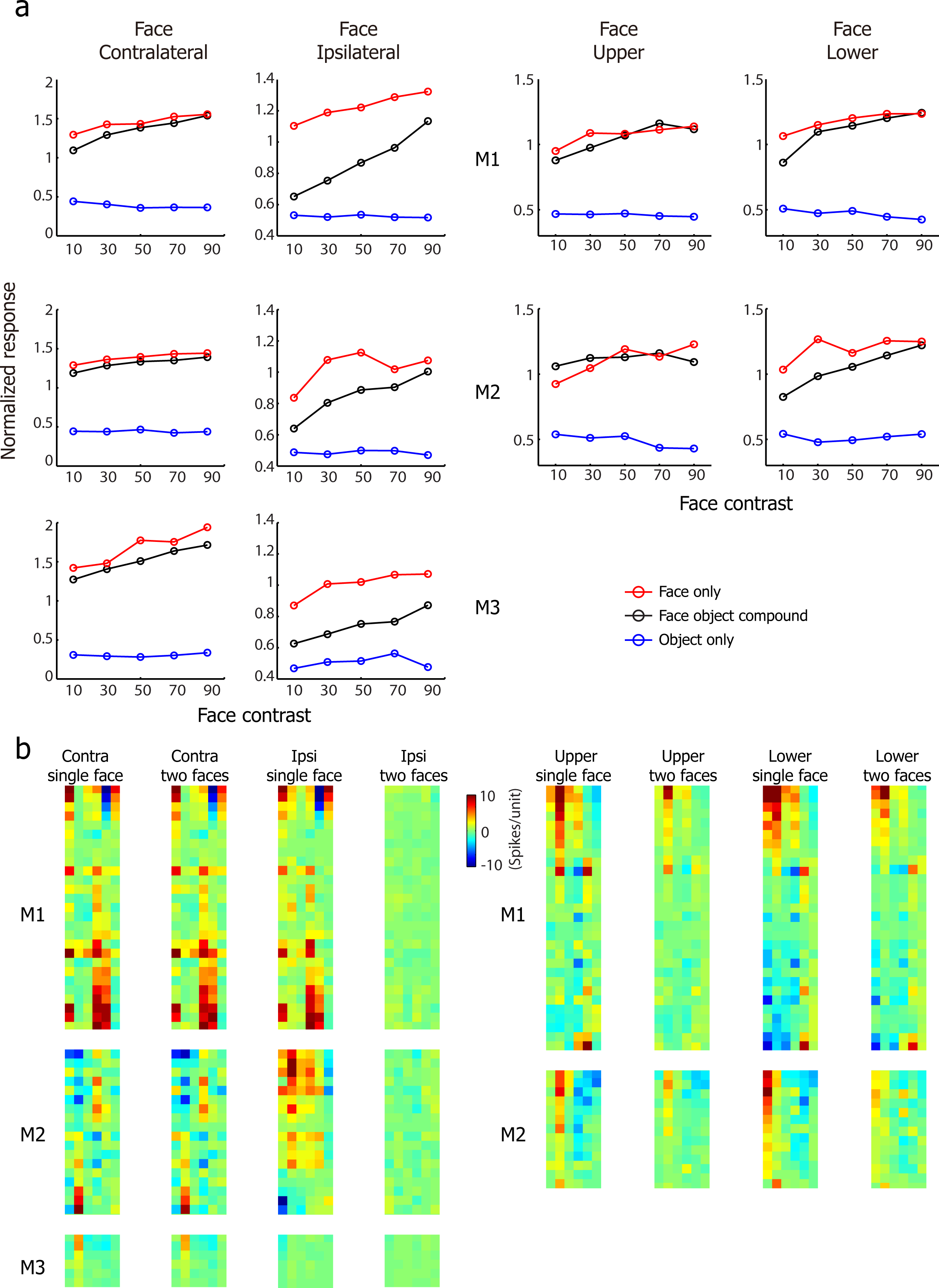
Main results shown separately for three monkeys. (**a**) Results of face-object experiment for monkeys M1, M2, M3. Conventions as in Figure 2a2, b2, c2, d2. (**b**) Results of face-face experiment for monkeys M1, M2, M3. Conventions as in Figure 4A2, B2.

**Figure S4.**
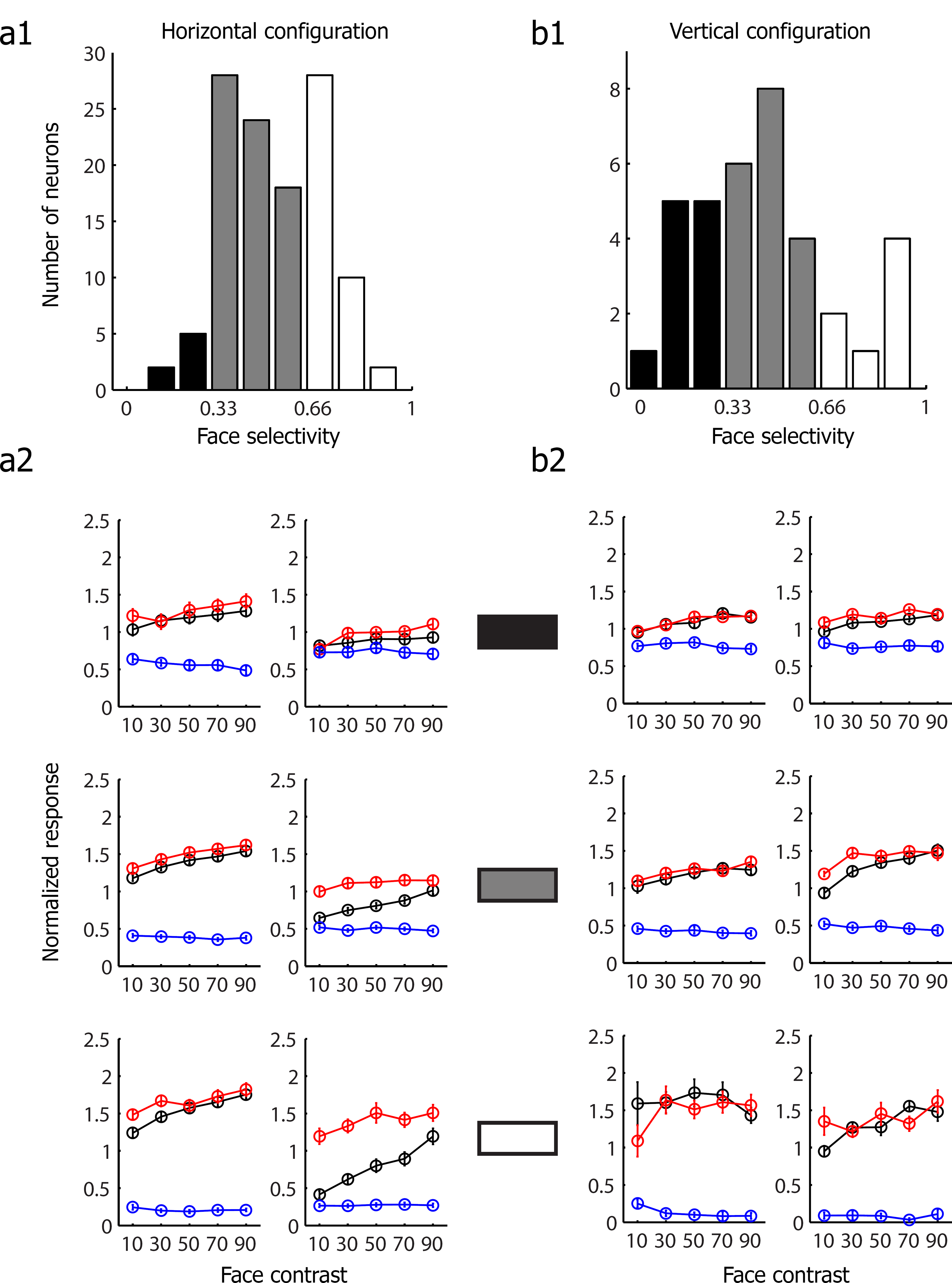
The integration rule for face-object pairs does not depend on a neuron’s face selectivity. (**a1**) The distribution of face selectivity, defined as FSI, across the population in the horizontal configuration. Neurons were classified into three groups: (1) low face selectivity (FSI < 0.33 black), (2) medium face selectivity (0.33 < FSI <0.66), (3) high face selectivity (FSI > 0.66, white). (**a2**) The population mean response of each group defined in Figure 2a2, b2. Conventions as in Figure 2 a1. (**b1**) Same as a1 for the vertical configuration. (**b2**) The population mean response of each group defined in Figure b1. Conventions as in Figure 2c2, d2.

**Figure S5.**
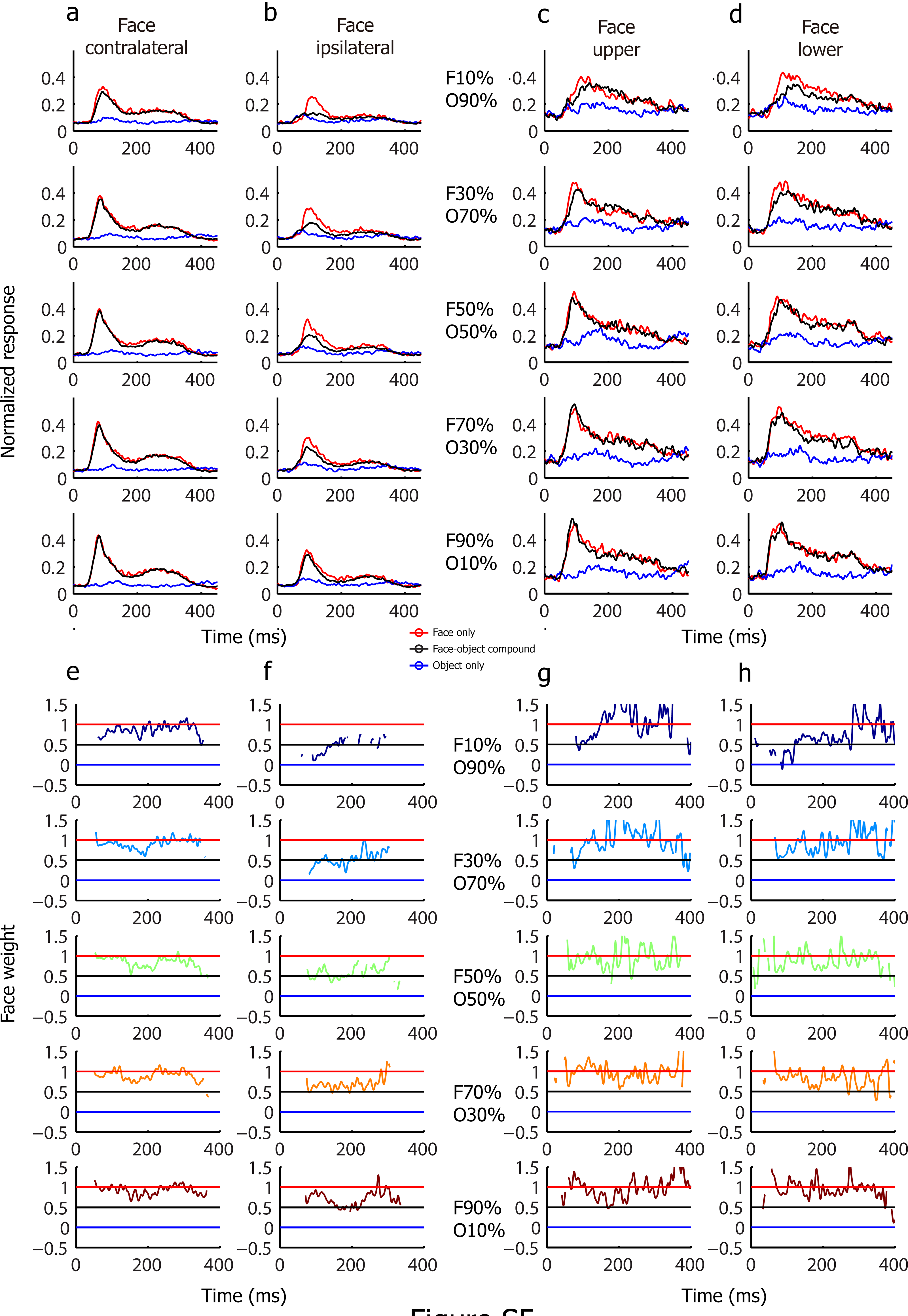
Response time course in response to face-object pairs. **(a)** The population mean response of neurons in ML to face-object pairs with face presented in the contralateral visual field and object presented in the ipsilateral visual field. Black lines represent the responses to the face-object pairs across time. Red (blue) lines represent the responses across time to the constituent face (object) of the corresponding face-object pair when presented in isolation. The 5 rows represent 5 different contrast levels. **(b)** Same as (a), for the condition in which faces were presented in the ipsilateral visual field and objects were presented in the contralateral visual field. **(c)** Same as **(a)**, for the condition in which faces were presented in the upper visual field and objects were presented in the lower visual field. **(d)** Same as (a), for the condition in which faces were presented in the lower visual field and objects were presented in the upper visual field. **(e)** The face weight (see Methods) computed across time at five relative contrast levels (from top to bottom) with face presented in the contralateral visual field and object presented in the ipsilateral visual field. Note some time points are missing because the difference between the responses to face and object was too small (<0.04), making the weight estimation meaningless. **(f)** Same as **(e)**, for the condition in which faces were presented in the ipsilateral visual field and objects were presented in the contralateral visual field. **(g)** Same as **(e)**, for the condition in which faces were presented in the upper visual field and objects were presented in the lower visual field. **(h)** Same as **(e)**, for the condition in which faces were presented in the lower visual field and objects were presented in the upper visual field.

**Figure S6.**
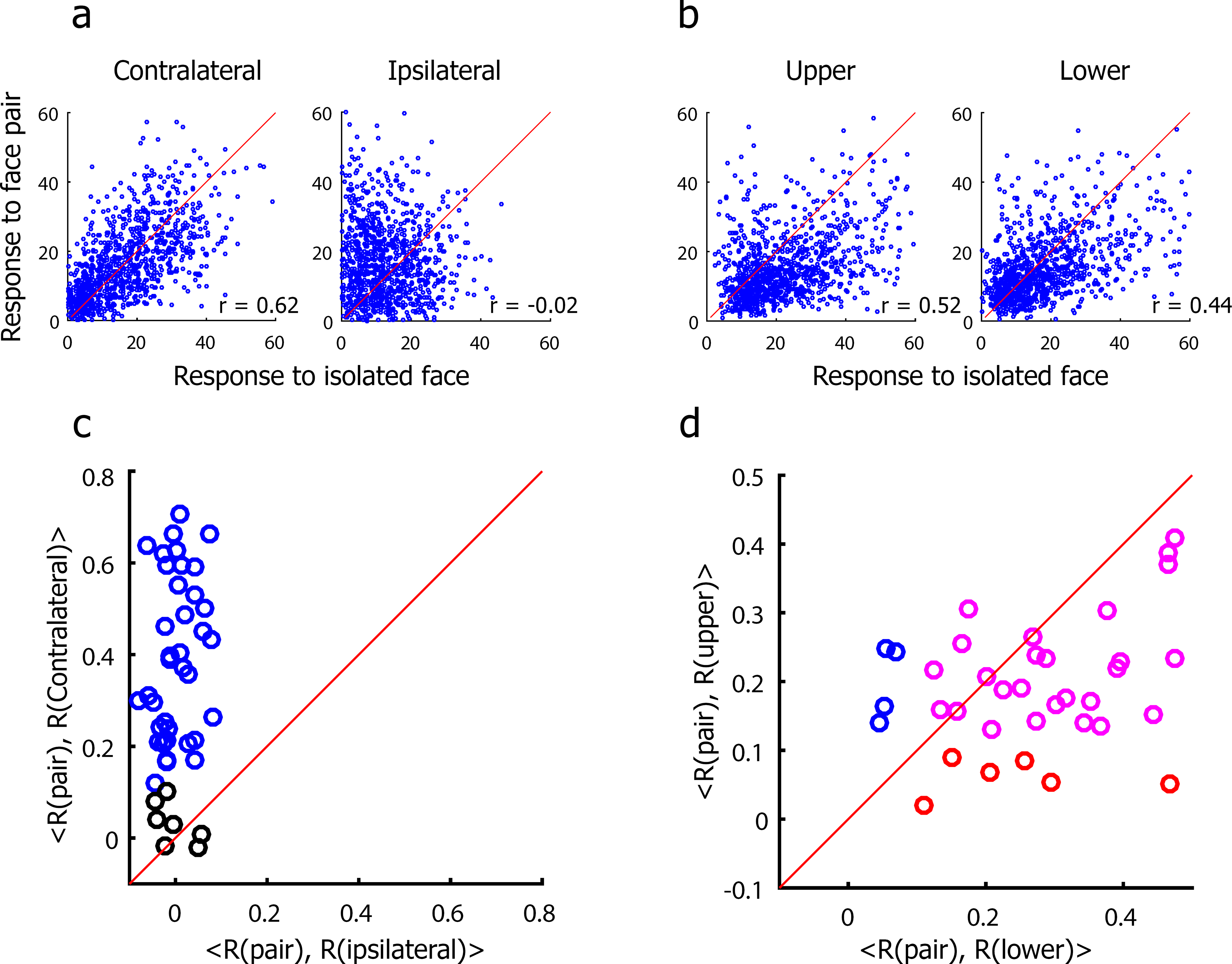
The correlation between response magnitudes to pairs of faces to isolated faces. (**a**) Response magnitudes to pairs of faces versus to isolated faces (left: contralateral face, right: ipsilateral face) for one example cell, with faces in the horizontal configuration. **(b)** Response magnitudes to pairs of faces versus to isolated faces (left: upper face, right: lower face) for one example cell, with face in the vertical configuration. **(c)** Scatter plots of the correlation between responses to pairs of faces and contralateral isolated faces versus the correlation between responses to pairs of faces and ipsilateral isolated faces, across all neurons. Blue indicates neurons that showed a significant correlation between the responses to pairs of faces and to contralateral isolated faces. Black indicates neurons that showed no significance for either correlation. (*p* < 0.05, Bonferroni corrected). **(d)** Scatter plots of the correlation between responses to pairs of faces and isolated faces above fixation versus the correlation between responses to pairs of faces and isolated faces below fixation across all the neurons. Blue indicates neurons that only showed a significant correlation between the responses to pairs of faces and isolated faces above the fixation. Red indicates neurons that only showed a significant correlation between responses to pairs of faces and isolated faces above fixation. Magenta indicates neurons that showed significance for both correlations (*p* < 0.05, Bonferroni corrected).

**Figure S7.**
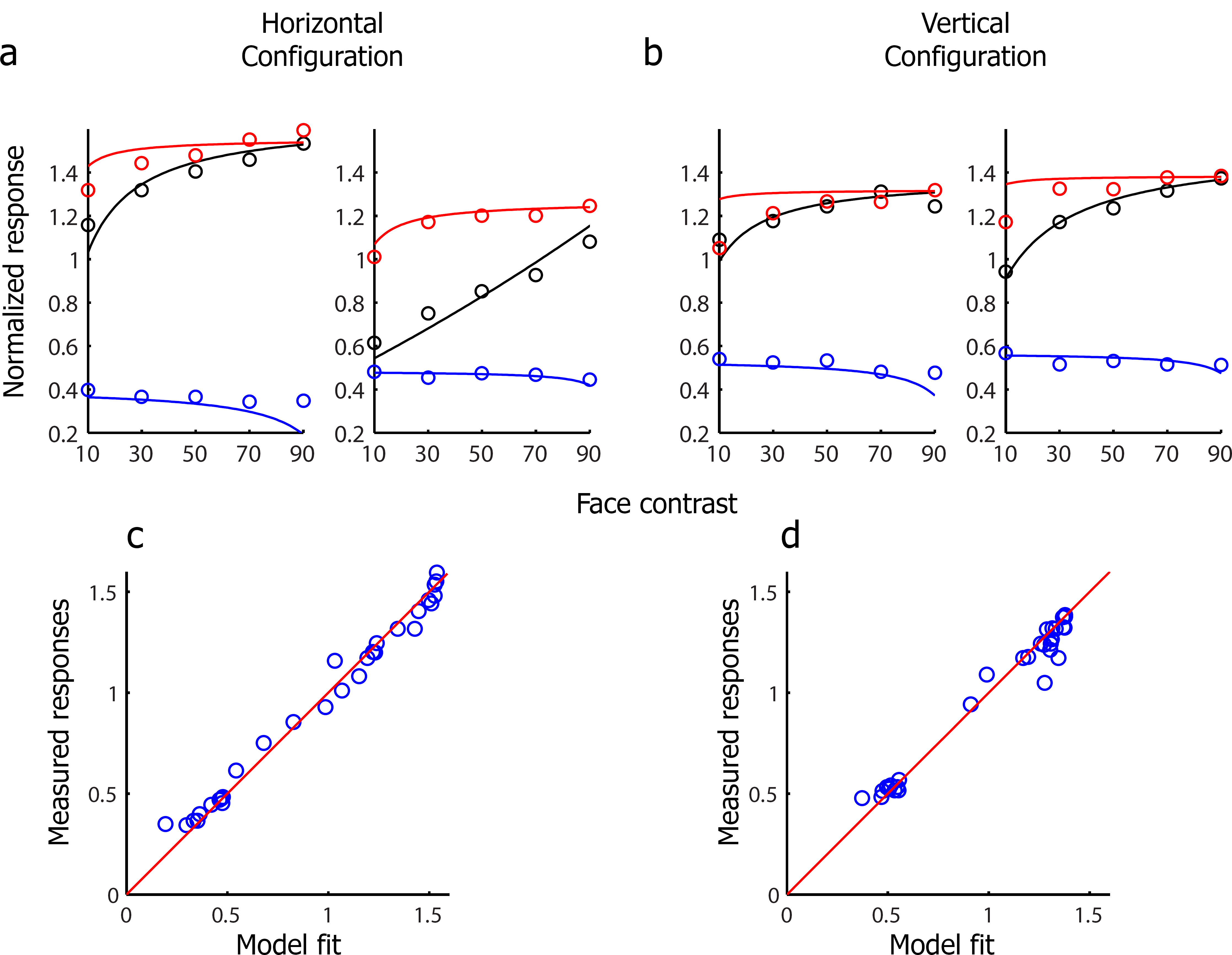
The normalization model using LFP amplitudes as weights predicts neural responses. **(a)** The normalization model fit using LFP amplitudes as weights to the face-object experiment when faces and objects were presented horizontally. Line indicates model fit, circles indicate observed data (data same as Figure 2a2, b2). **(b)** Same as (a), for faces and objects presented vertically. **(c)** The correlation between the observed responses and the normalization model fit when faces and objects were presented horizontally. We added a free LFP baseline parameter to obtain the model fits (i.e., the weight in the normalization model were calculated as the LFP amplitude minus the baseline). **(d)** Same as (c), for face and objects presented vertically.

**Figure S8.**
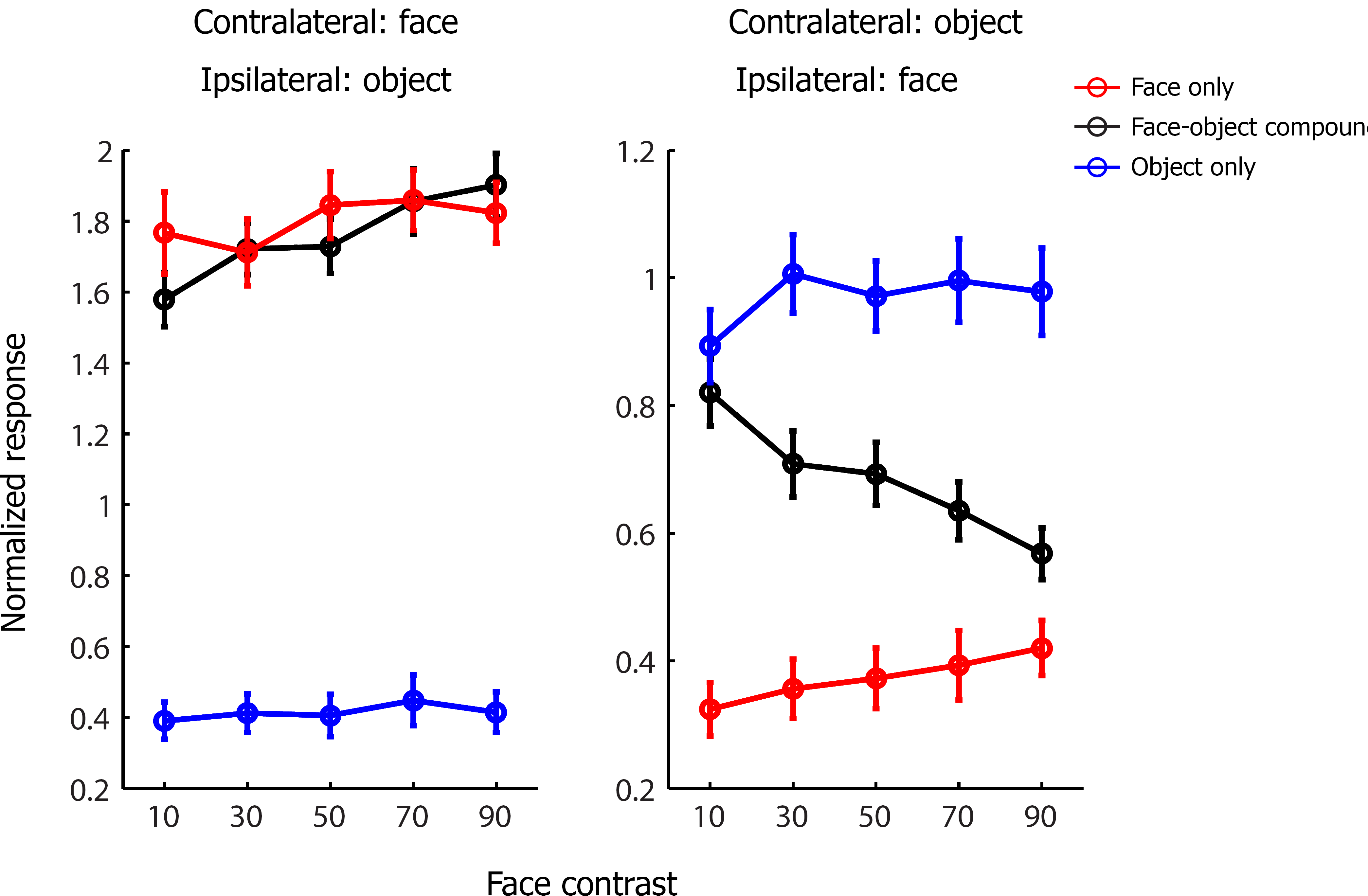
Integration rule for subset of face cells showing a greater response to contralateral objects compared to ipsilateral faces. Black line represents response to face-object pair at different relative contrasts. Red (blue) line represents responses to the constituent face (object) of the corresponding face-object pair when presented in isolation.

**Figure S9.**
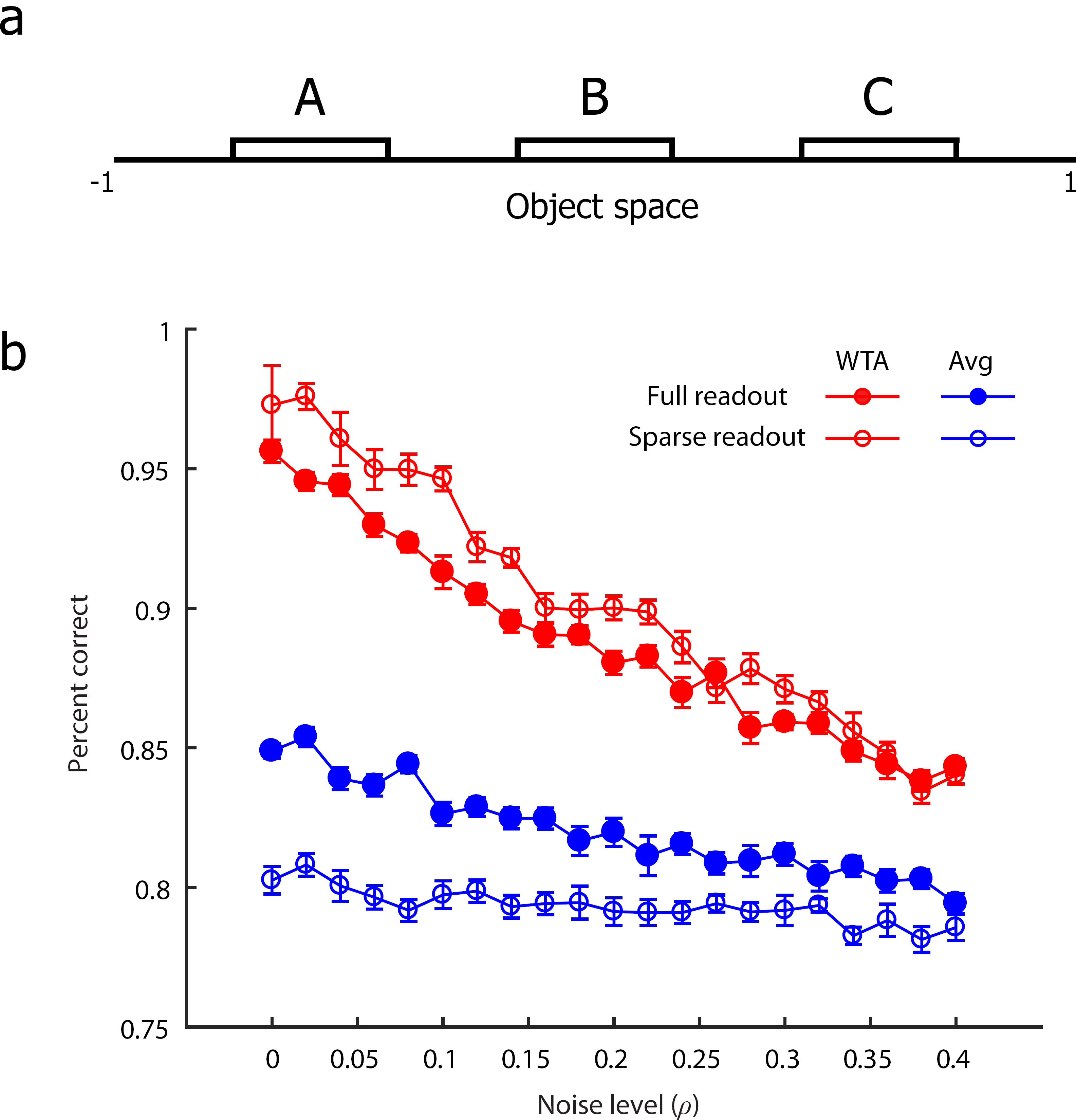
Simulation of object decoding as a function of integration rule (winner-take-all, averaging), readout strategy (sparse, full), and noise level. (**a**) Hypothetical one dimensional object space. (**b**) Decoding accuracy (averaged across three classifiers, for A/not A, B/not B C/not C) as a function of noise level, for a winner-take-all rule (red curves) and averaging rule (blue curves). Filled circles indicate performance when reading out all neurons, while open circles indicate performance when reading out neurons selective for the object being detected. The winner-take-all rule always outperformed the averaging rule, with especially pronounced difference for sparse readout.

**Figure S10.**
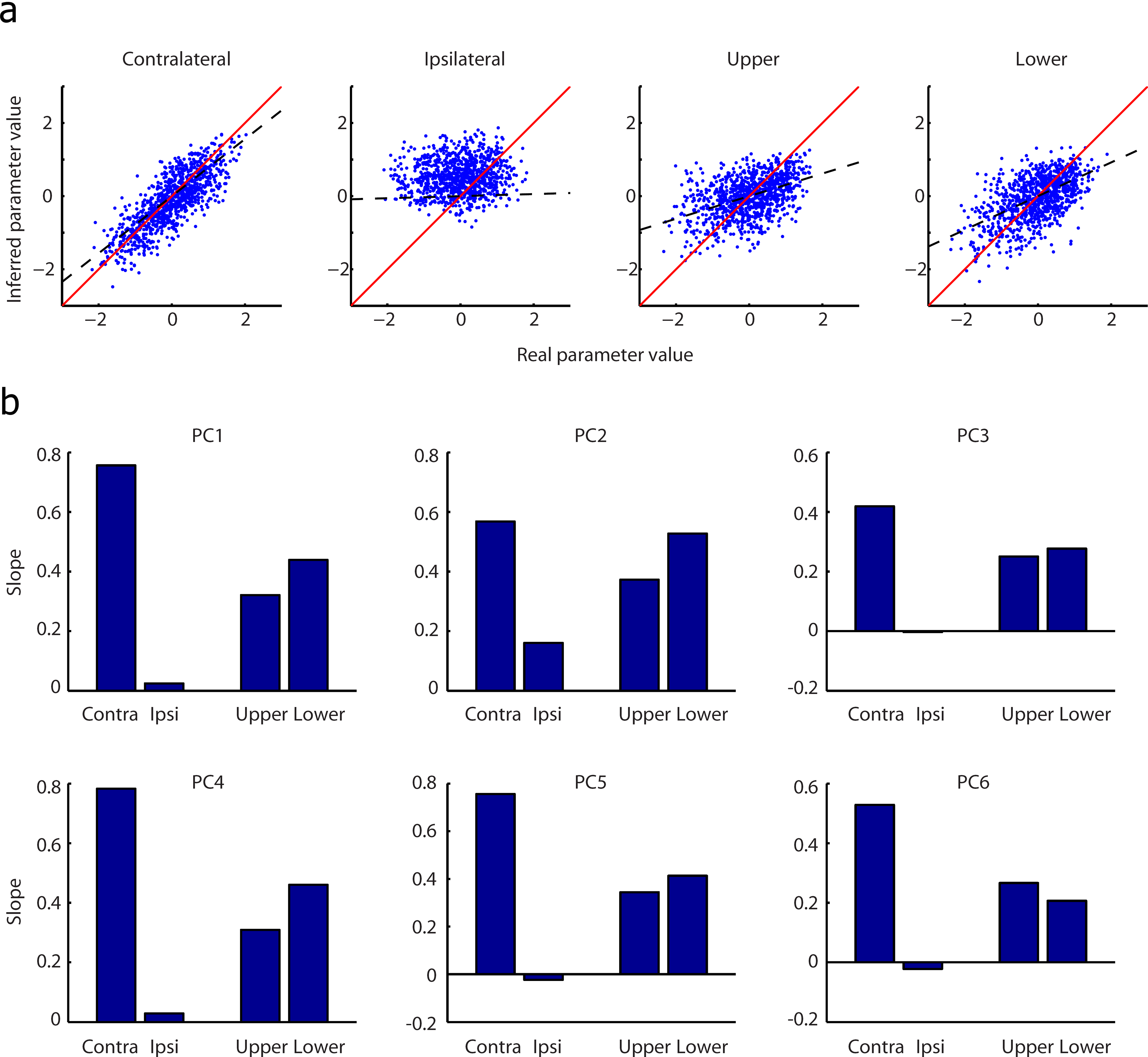
Decoding face features in pairs of faces across different spatial configurations. (**a**) For PC1 (i.e., dimension 1 of our 6-d face space), we plot the predicted versus actual feature value, for a face presented contralaterally or ipsilaterally in a horizontal pair, and for a face presented above or below in a vertical pair. The predicted values were determined according to the decoding model in Supplementary Text 2. (**b**) The slope of the best-fit line between predicted and actual feature values for each of the 6 dimensions in our 6-d face space, for all four spatial conditions.

**Author Contributions:** Conceptualization, P.B. and D.Y.T.; Methodology P.B. and D.Y.T.; Investigation, P.B.; Writing – Original Draft P.B. and D.Y.T.; Writing – Review & Editing, P.B. and D.Y.T.; Funding Acquisition D.Y.T.

